# Dissecting heterogeneity in cortical thickness abnormalities in major depressive disorder: a large-scale ENIGMA MDD normative modelling study

**DOI:** 10.1101/2025.03.17.643677

**Authors:** J.M.M Bayer, L.S van Velzen, E Pozzi, C Davey, L.K.M Han, S.E.E.C Bauduin, J Bauer, F Benedetti, K Berger, L.M Bonnekoh, K Brosch, R Bülow, B Couvy-Duchesne, K.R Cullen, U Dannlowski, D Dima, K Dohm, J.W Evans, C.H.Y Fu, P Fuentes-Claramonte, B.R Godlewska, J Goltermann, A Gonul, R Goya-Maldonado, H.J Grabe, N.A Groenewold, D Grotegerd, O Gruber, T Hahn, G.B Hall, J Hamilton, B.J Harrison, S.N Hatton, M Hermesdorf, I.B Hickie, T.C Ho, N Jahanshad, A Jansen, A.J Jamieson, T Kamishikiryo, T Kircher, B Klimes-Dougan, B Krämer, A Kraus, A Krug, E.J Leehr, R Leenings, M Li, A McIntosh, S.E Medland, S Meinert, E Melloni, B Mwangi, I Nenadić, G Okada, M. Oudega, M.J Portella, E Rodríguez, L Romaniuk, P.G. Rosa, M.D Sacchet, R Salvador, P.G Sämann, H Shinzato, K Sim, E Simulionyte, J.C Soares, D.J Stein, F Stein, A Stolicyn, B Straube, L.T Strike, L Teutenberg, F Thomas-Odenthal, S.I Thomopoulos, P Usemann, N.J.A van der Wee, H Völzke, M. Wagenmakers, M Walter, H.C Whalley, S Whittle, N.R Winter, K Wittfeld, M Wu, T.T Yang, C.A Zarate, G.B Zunta-Soares, P.M Thompson, D.J Veltman, A.F Marquand, L Schmaal

## Abstract

**Importance:** Major depressive disorder (MDD) is highly heterogeneous, with marked individual differences in clinical presentation and neurobiology, which may obscure identification of structural brain abnormalities in MDD. To explore this, we used normative modeling to index regional patterns of variability in cortical thickness (CT) across individual patients.

**Objective:** To use normative modeling in a large dataset from the ENIGMA MDD consortium to obtain individualised CT deviations from the norm (relative to age, sex and site) and examine the relationship between these deviations and clinical characteristics.

**Design, setting, and participants:** A normative model adjusting for age, sex and site effects was trained on 35 CT measures from FreeSurfer parcellation of 3,181 healthy controls (HC) from 34 sites (40 scanners). Individualised z-score deviations from this norm for each CT measure were calculated for a test set of 2,119 HC and 3,645 individuals with MDD. For each individual, each CT z-score was classified as being within the normal range (95% of individuals) or within the extreme range (2.5% of individuals with the thinnest or thickest cortices).

**Main outcome measures:** Z-score deviations of CT measures of MDD individuals as estimated from a normative model based on HC.

**Results:** Z-score distributions of CT measures were largely overlapping between MDD and HC (minimum 92%, range 92-98%), with overall thinner cortices in MDD. 34.5% of MDD individuals, and 30% of HC individuals, showed an extreme deviation in at least one region, and these deviations were widely distributed across the brain. There was high heterogeneity in the spatial location of CT deviations across individuals with MDD: a maximum of 12% of individuals with MDD showed an extreme deviation in the same location. Extreme negative CT deviations were associated with having an earlier onset of depression and more severe depressive symptoms in the MDD group, and with higher BMI across MDD and HC groups. Extreme positive deviations were associated with being remitted, of not taking antidepressants and less severe symptoms.

**Conclusions and relevance:** Our study illustrates a large heterogeneity in the spatial location of CT abnormalities across patients with MDD and confirms a substantial overlap of CT measures with HC. We also demonstrate that individualised extreme deviations can identify protective factors and individuals with a more severe clinical picture.

**Key points:** 

**Question:** Can z-scores derived from normative modelling shed light on the heterogeneous group-level findings of cortical thickness abnormalities in major depression and what characterises individuals at the extreme ends of cortical thickness abnormalities?

**Finding:** We confirmed a large overlap in z-score distributions between depressed individuals and healthy controls and a heterogeneous spatial distribution of extreme z-deviations across brain regions across individual patients. Lower z-scores for cortical thickness were related to more severe clinical characteristics.

**Meaning:** Our findings confirm the heterogeneity in individual variation in the location and extent of CT abnormalities across patients with MDD and stress the importance of individualised predictions when examining cortical thickness abnormalities.

## 1. Introduction

Major depressive disorder (MDD) is a highly prevalent mental illness, impacting more than 300 million people worldwide^1–3^. Neuroimaging research in the last decades has aimed to identify the neural basis of MDD. Cortical thickness has shown to be genetically and phenotypically independent from other imaging phenotypes and to be affected by complex and distinct pruning and myelination processes related to typical and atypical learning and development trajectories in clinical and subclinical populations^4–8^. While earlier studies found medium to large effect sizes for cortical thickness (CT) alterations in MDD (Cohen’s *d* 0.48-0.60)^9,10^, more recent studies on CT suggest that these effect sizes have been overestimated, potentially due to small sample sizes and publication bias (e.g., maximum Cohen’s d=0.13, in larger samples^11–13^). Rather, having a diagnosis of MDD seems to only subtly affect regional CT, evidently in frontal (including orbitofrontal cortex (OFC) areas, the anterior cingulate (ACC), the dorsomedial prefrontal cortex (PFC)) and posterior cingulate and temporal areas^12,14,15^. These small observed effect sizes imply that many individuals with MDD show CT values that overlap with those of healthy individuals^11,16^, and thus cannot be directly used for classification purposes.^17^

The association between these small effect sizes and the large inter-individual clinical heterogeneity within MDD^18–20^ remains unresolved. To a certain degree, stratifications of MDD by clinical features (such as age of onset or use of antidepressants) point to subgroups with stronger morphological alterations^12,21,22^. Still, there remains a large portion of MDD patients with CT patterns practically not distinguishable from HC. In addition, while most studies find thinner cortices^23–25^ to be associated with depression, also thicker cortices^24,26^, or no difference^27^ have been reported, especially in periods of rapid brain maturation such as adolescence^28^. Due to the general change of CT with age^29,30^, these CT findings may be the result of a complex interaction between the effects of maturation, ageing and the MDD diagnosis.

The group-average comparison approach that is typical of neuroimaging studies is meant to look at the group as a whole and does not identify individual differences within a group, as it treats individualised variability as a nuisance, allowing only for inferences at the group level. In addition, a complex interaction between brain maturation or ageing and diagnosis may be obscured by limiting the analysis to group comparisons, particularly when age-interaction effects are not modelled. Normative modelling^31,32^ (NM) is an alternative approach to identifying morphological brain alterations, with a focus on positioning each individual’s brain measure on a normative scale. The framework has been successfully applied to clinical neuroimaging data including CT measures, uncovering the anatomical heterogeneity in various mental disorders^28,29,33–39^. NM establishes the range of (statistically) normal variation in the data based on the covariates in the model (e.g., age and sex) and subsequently parses each individual’s raw score onto an individualised z-score deviation from that norm. As one practical use case, NM allows to detect individuals with extreme deviations (e.g., +/- 2 standard deviations) from the norm, which might be beneficial in cases of large heterogeneity, subtypes^40^, or where anatomical differences are subtle^12^.

Beyond this, multi-site neuroimaging data are sensitive to effects induced by site differences^41–43^. Especially in the case of small effect sizes, site effects can thus easily overshadow effects of interest. Extending previous work^44^, we have developed linear and non-linear versions of normative models that can deal with these site differences^43^. Our approach thus combines the individualization aspect of NM with the ability to correct for site effects, making normative modelling applicable for multi-site neuroimaging data^43,44^.

In this study, we aim to elucidate the heterogeneity in CT alterations across individuals with MDD by examining the variation in CT z-scores for different brain regions in a large, pooled depression data set from the ENIGMA MDD consortium using NM^45,46^. First, we aim to re-estimate group average differences in CT between MDD and HC based on z-scores from NM that reflect an age, sex and site normed metric, hypothesizing a similar anatomical CT deficit pattern compared to our previous ENIGMA MDD findings of CT alterations in MDD^12^. Further, we aim to investigate characteristics of individuals with extreme deviations in either direction (positive and negative deviations), quantifying their frequency and anatomical distribution. For this we explored the association of extreme CT deviations with clinical and lifestyle features, including symptom severity, number of depressive episodes, use of antidepressant medication, age of onset, childhood trauma and body mass index (BMI).

## 2. Methods

### 2.1. Sample

The ENIGMA MDD working group combines neuroimaging and clinical data across 4,597 MDD patients and 5,926 healthy controls (HC) from 53 international cohorts. Data from 34 cohorts were included in this study. All sites obtained approval from local institutional review boards and ethics committees. All participants provided informed consent.

### 2.2. Demographic and clinical characteristics

Demographic information include age, sex, BMI and MDD status (HC, or MDD; for demographics, clinical characteristics, in- and exclusion criteria, diagnostic instruments used and scan protocol details per cohort, see **Tables S1.1-3**). Clinical information included episode status (HC, first episode depression, recurrent episode depression), remission status (HC, acutely depressed, remitted), antidepressant use at the time of scan (HC, antidepressant-free, antidepressant user), age of onset of depression (MDD with =< 21 years age of first onset, MDD with > 21 years age of onset^12,21^), Beck Depression Inventory (BDI)^47^ total score (only for MDD), Hamilton Depression Rating Scale^48^ (HDRS, only for MDD) total score and Childhood Trauma Questionnaire (CTQ) total score^49^ (see **Tables S1.4-5**).

### 2.3. Image processing and analysis

3D structural T1-weighted brain MRI scans were acquired at each site (image acquisition parameters are detailed in **Table S1.3**). Analysis and quality control of the images were performed locally using harmonised protocols provided by the ENIGMA consortium (http://enigma.ini.usc.edu/protocols/imaging-protocols). Cortical parcellations were acquired using FreeSurfer (versions 5.1 and 5.3)^50^ based on the Desikan-Killiany atlas^51^ (34 regions in the left and 34 in the right hemisphere, plus one average across all regions). Measures were visually inspected and statistically analysed for outliers using standardised ENIGMA protocols http://enigma.ini.usc.edu/protocols/imaging-protocols). As our prior studies did not show lateralisation effects of CT alterations in MDD^52^, CT values were averaged for each region across the left and right hemispheres, resulting in 34 bilateral regions and one whole-brain average CT measure. The final sample included in this study amounted to 5,300 HCs and 3,645 individuals with MDD from 34 sites including 40 different MR scanning platforms.

### 2.4. Training and test set

HC data was divided into a training set (60% of HCs, N=3,181) and test set (40% of HCs, N= 2,119) using a within-site split. The normative model was not trained on all HCs to ensure that potential differences between HC and MDD in deviation z-scores were not due to worse fit of the model in the MDD group compared to HC because the model was trained on all HCs. The training and test sets had comparable distributions in terms of age and sex (**Figure 1**, for a more detailed analysis see **supplemental text S2.2a and table S2.2a,b**). All individuals with MDD (N=3,645) were part of the MDD test set. The MDD test and the HC test sets were imputed, scaled, and centred across sites, based on the HC training set using k=10 nearest neighbour imputation. An overview of the data processing pipeline may be found in **Figure 1** and **supplemental S2.2.**

**Figure 1.**
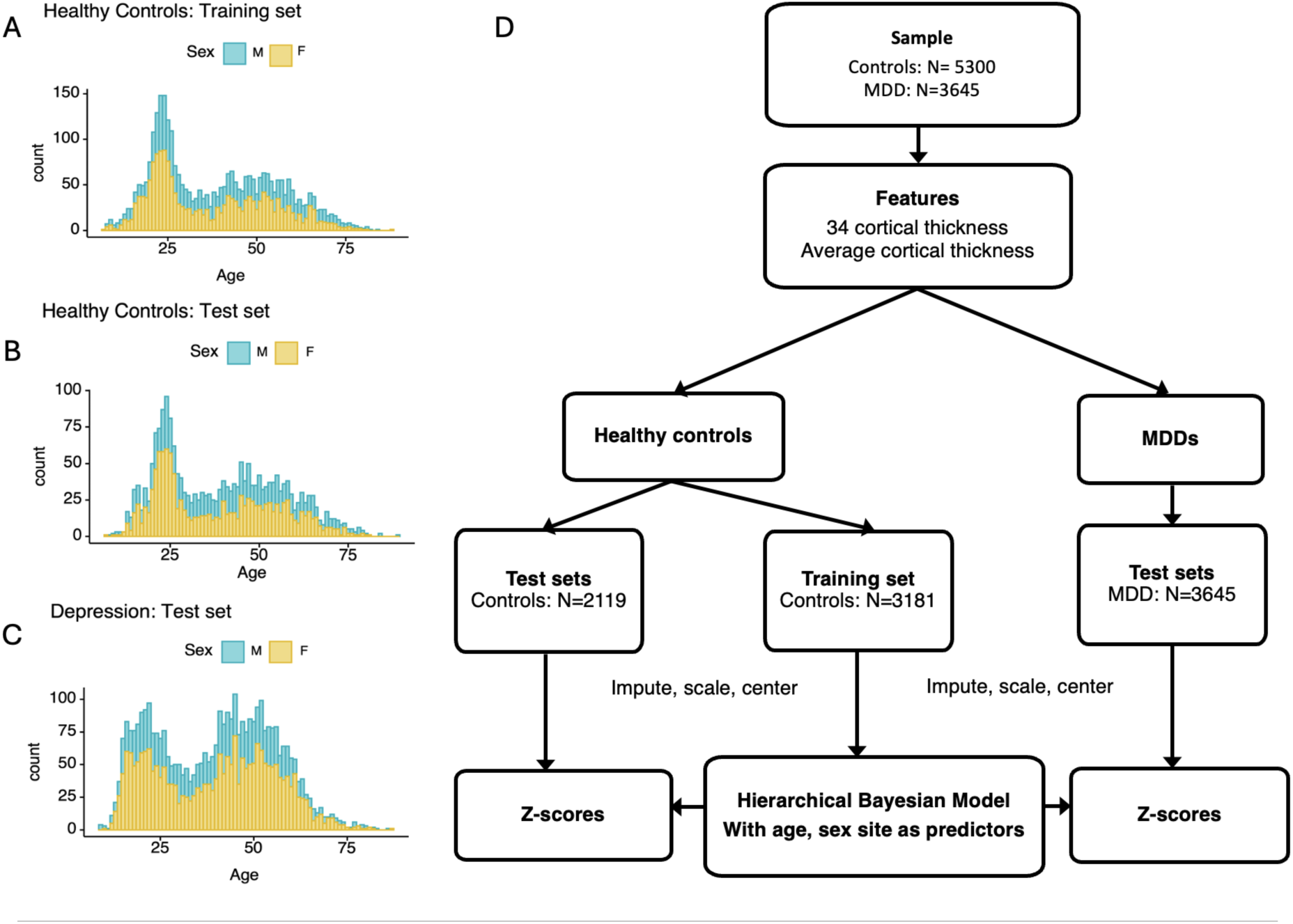
Distribution of age and sex for the three partitions of the data used in this study. A) Training set, healthy controls. B) Test set, healthy controls. M= male, F=female C) Test set, MDD. D) Illustrated Pre-Processing pipeline. The sample consisted of 34 regional features and one whole brain measure from 5300 healthy controls and 3645 individuals with MDD from the ENIGMA MDD consortium.

### 2.5. Normative Modelling

We applied our previously developed NM framework predicting CT measures based on age, sex and site^43^. A hierarchical Bayesian approach in *Stan,* described elsewhere^43^, was applied, including the covariates, age, sex and site. A more in-depth description of the NM process may be found in **supplemental text S2.4**. All code can be found in: https://github.com/likeajumprope/Normative-ENIGMA-MDD. An interactive version of all figures can be found here: https://likeajumprope.github.io/Normative-ENIGMA-MDD/

### 2.6. Statistical analysis

All statistical analyses reported below were performed on deviations in CT from the norm (z-scores) derived from the MDD and HC test sets. A hierarchical Bayesian regression model with age, sex and site as predictors was used to create a model mapping the normative variation in each CT measure onto age and sex, while accommodating for multi-site effects^43^. We further used a B-splines model with 5 knots to model non-linear associations between age and CT, which allowed us to calculate z-scores for each participant *i* and region *j* using the following:

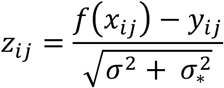

Where *f*(*x_ij_*) is the prediction of the model, *y_ij_* is the true value, σ^2^ is a random gaussian noise component and σ^2^ is the predictive variance from the model (for more details see ^43,53^).

#### 2.6.1. Average group differences in z-scores per ROI

T-tests were performed to examine MDD vs HC group differences in z-scores. Region-wise overlap scores, Cohen’s *d* metrics and distributional overlap percentages between HCs and individuals with MDD^11^ were also calculated. Due to residual associations with z-scores, age and sex were included as covariates in all statistical models described below. P-values were corrected using false-discovery-rate (FDR) correction for 35 statistical models (34 regions + 1 global brain measure).

#### 2.6.2. Extreme deviations per ROI and global measure

Individuals were divided into subgroups for each bilateral region, based on their z-score deviation: (1) falling within the *norm* (*normal*; between and 2.5^th^ and 97.5^th^ percentile, or z-scores between -1.96 or 1.96), (2) *extreme positive deviators* from the norm (*supra-normal*; above 97.5th^th^ percentile with z-scores >= 1.96), and (3) *extreme negative deviators* (*infra-normal*; below 2.5^th^ percentile, with z-scores =< -1.96). This resulted in a 0 (normal), -1 (infra-normal) or 1 (supra-normal) score for each cortical region for each individual. Percentages of individuals with MDD with at least one infra- or supra-normal z-score were compared to the percentages of HCs with infra- and supra-normal z-scores using a chi-squared test.

#### 2.6.3. Spatial location indifferent summary deviation scores

To account for the heterogeneity in the spatial location of the infra- and supra-normal z-scores across different individuals, we derived the following spatial location indifferent summary measures that target the analysis of extreme deviations^54,55^:

- *Average z-score*: the average z-score across all 34 cortical regions.
- *Load score*: individual total number of regions with infra- or supra-normal z-scores across all brain regions. Load scores were created by separately summing the number of infra and supra normal z-score deviations per individual, resulting in one positive and one negative load scores with a range from 0 – 35 each^54,55^.
- *Extremity score*: z-score with the largest negative or positive value across all regions per individual resulting in one positive and one negative extremity score per individual^54,55^.

## 3. Results

### 3.1. Model fit

Model fits of the normative models to the training were satisfactory in terms of Explained Variance, Root Mean Squared Error, Correlation and Mean Standardized Log Loss for the test sets. A distribution of extreme deviations close to the expected values of 5% for both test sets indicated a good generalisability of the training model **(Table S1.9)**. A summary can be found in **supplemental text S2.5** and in **tables S1.10-13**.

### 3.2. Group average differences in z-scores

Significantly lower z-scores in individuals with MDD compared to HC were observed in 25 out of 35 brain regions, with greatest differences in the fusiform gyrus (Cohen’s *d*=-0.17), inferior (*d*=-0.13) and middle temporal gyrus (*d*=-0.13), the bank of the superior temporal sulcus (*d*=-0.12) and the insula (*d*=-0.12, **Figure 2d,e**). These results aligned with those of a previous publication^12^ (*ρ*=0.70, see **supplemental text S2.6**). No average regional CT increases in MDD compared with HC were detected. Across all regions, z-score distributions were shifted towards more negative in MDD (**Figure 2f**). The percentage of z-score distribution overlap between HC and MDD ranged between 92% and 98%. The fusiform gyrus, the inferior frontal gyrus pars opercularis and average CT showed the lowest overlap, indicating the largest difference in z-score distributions between HC and MDD (92.3%, p<0.001; 93.3%, p<0.001; 93.4% p<0.001; respectively; **Figure 2a-c, Table S1.7)**. For a full region-wise overview of Cohen’s *d* and distributional overlap scores, see **Table S1.7**.

**Figure 2.**
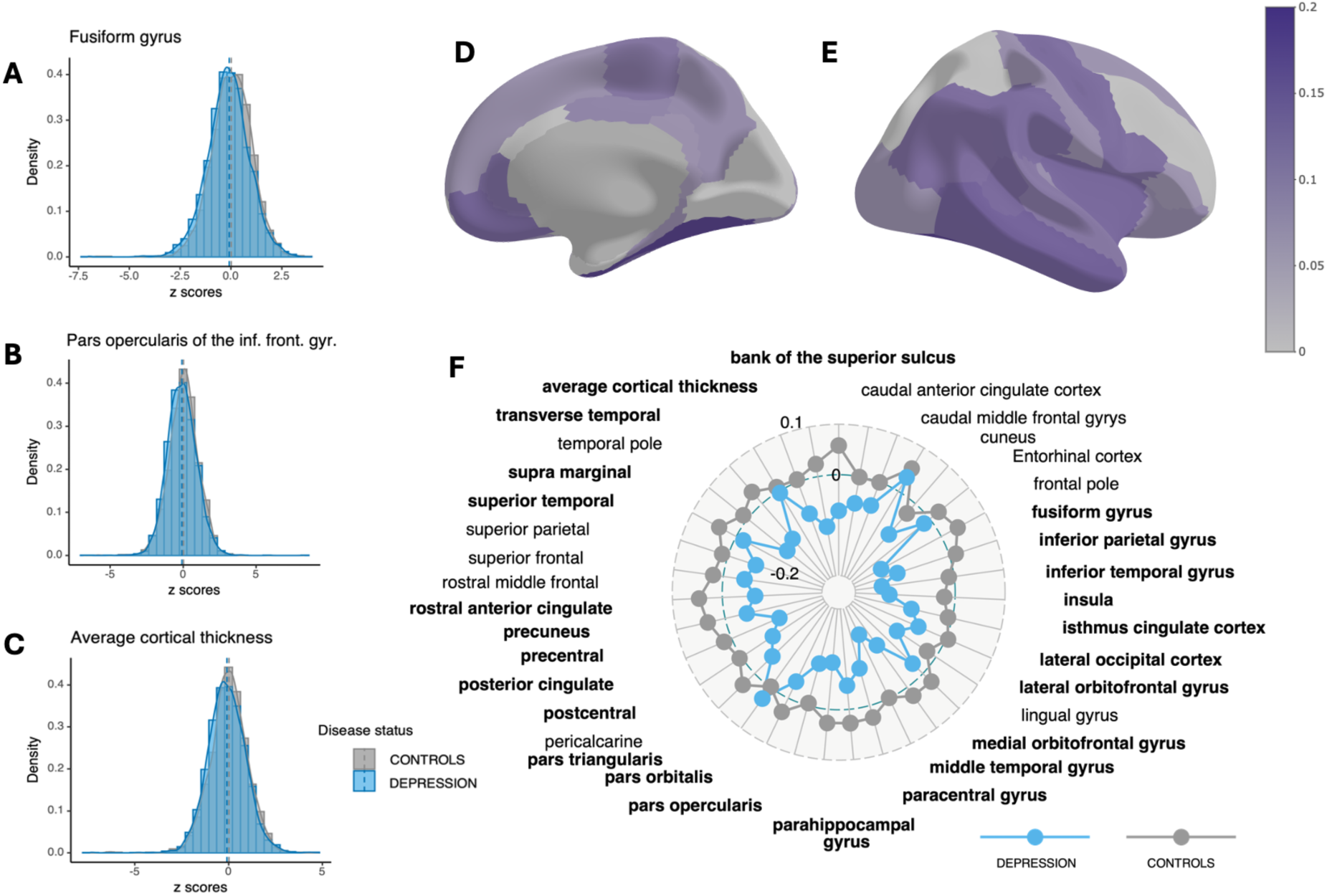
**A, B, C:** Distributional overlap between individuals with MDD and HCs (HC) for the A. fusiform gyrus, B: the pars opercularis of the inferior frontal gyrus and C: average CT, showing the lowest distributional overlap between MDD and HC groups. **D, E:** Cohen’s d effect sizes of differences in z-scores between MDD and HCs for each significant cortical region (FDR corrected, only significant regions are displayed). Red: Individuals with MDD have lower z-scores (thinner cortices) on average than HCs. An overview of Cohen’s d effect sizes for z-score differences in different cortical regions is provided in Table S1.7 in the supplementary material. D) medial view. E) lateral view. **F:** Spider plot showing the average z-scores for individuals with MDD and HCs for all cortical regions. All regions that show significant differences are plotted in bold.

### 3.3. Heterogeneity in spatial location of extreme negative and positive deviations

In MDD, 30.6% of individuals showed at least one cortical region with a supra-normal z-score (extreme positive deviation), versus 32.1% in HC (*χ*^2^=11.755, p<0.001), while 34.5% of individuals with MDD had at least one cortical region with an infra-normal z-score (extreme negative deviation) compared to 30.0% of HC (*χ*^2^=12.305, p<0.001). An illustration of the number of regions with extreme deviations is given in **Figure 3**.

**Figure 3.**
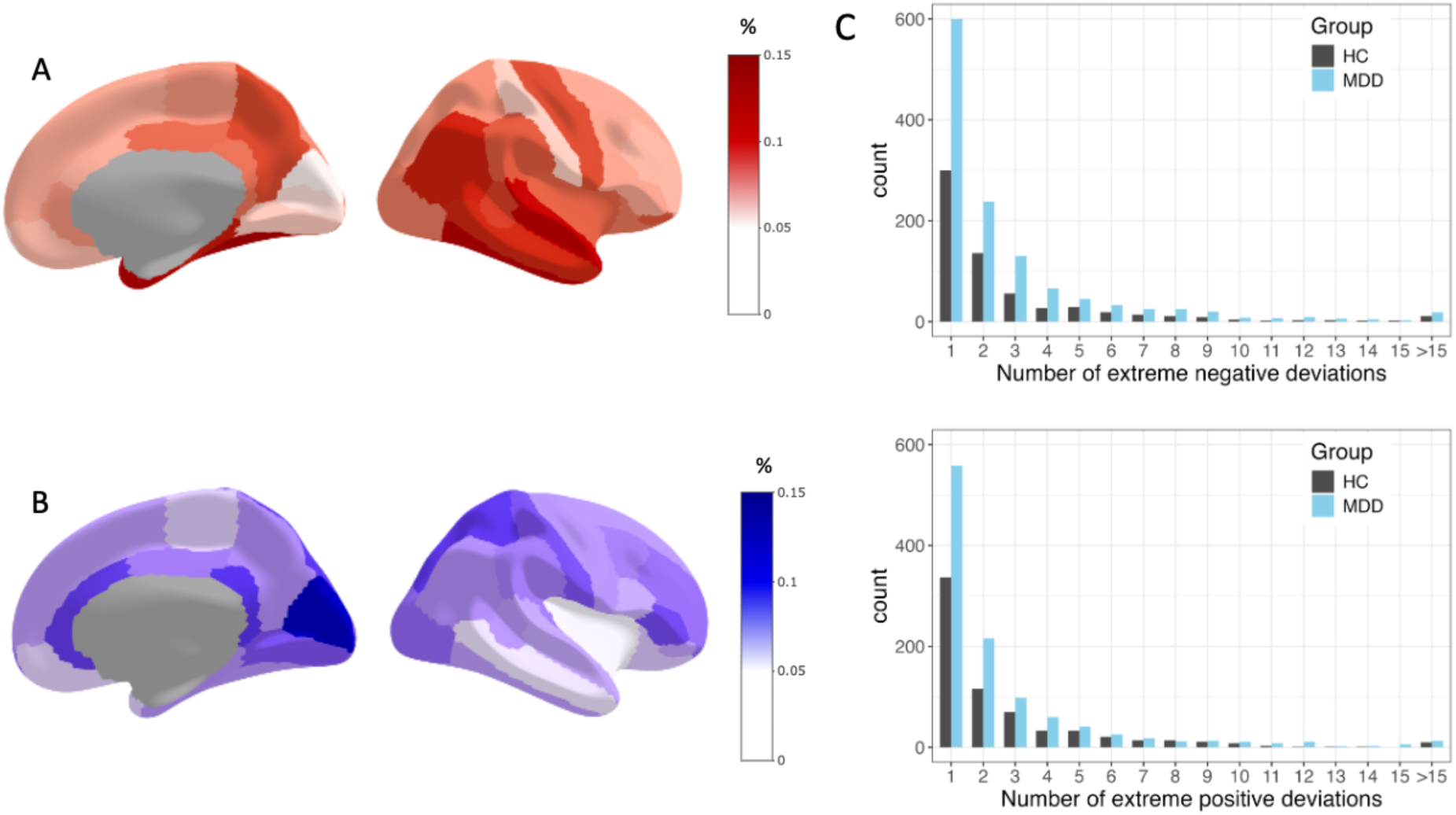
**A. B:** Heterogeneity in cortical thickness deviations in depression. Percentage of individuals with MDD with an infra-normal z-scores (extreme negative deviations; z < -1.96) and supra-normal z-scores (extreme positive deviations; z >1.96) per cortical region. **(A)** infra-normal z-scores; **(B)** supra-normal z-scores. **C:** Distribution of positive load score (number of extreme negative deviation; z > 1.96) **(C, top)** and negative load score (number of extreme positive deviations; z < -1.96) **(C, bottom)** in individuals with MDD and healthy controls.

No more than 12.1% (pericalcarine gyrus, cuneus) of individuals with MDD showed a positive extreme (supra-normal) z-score within the same region (**Figure 3b**) and no more than 11.9% (fusiform gyrus) of individuals showed an infra-normal z-score within the same region (**Figure 3a**). This illustrates the heterogeneity that is evident in the spatial location of extreme negative and positive deviations across individuals with MDD. A list of the percentages of extreme deviations for each region can be found in **Table S1.8**. Example profiles of individuals with the highest negative load and extremity score and the most negative average z-score can be found in **S2.10**.

### 3.4. Deviation scores and correlation with clinical variables

An overview of all associations between all summary deviation scores and clinical covariates is given in **Table 1** and in **Tables S1.14-S1.18 and S2.9**.

**Table 1.** Odds ratios and effect direction from predicting clinical covariates with brain summary scores with logistic and linear regression. Reference category in *italics*. MDD: Major depressive disorder. HC: healthy controls. FE: First episode depression. RE recurrent episode depression. ACUTE: acutely depressed. REM: remitted depressed. AD use: antidepressant use at the time of the scan. AD free: antidepressant free at the time of the scan. EAO: early age of onset (=< 21 years). LAO: late age of onset (> 21 years). BDI: Beck depression inventory. HDRS Hamilton depression rating scale. BMI: Body Mass Index. * main effects reported, interaction effect with MDD status not significant. **Neither main nor interaction effects significant. ↑ increase. ↓ decrease. *pt*: point. FDR adjusted. n.s. means no significant association was found. Full tables can be found in supplementary material S1.14-S1.18.

| Independent variable |  | Average z ↑ | Positive Load ↑ | (absolute) Negative Load ↑ | Positive extremity ↑ | (absolute) Negative extremity ↑ |
| --- | --- | --- | --- | --- | --- | --- |
| Dependent var. |  |  |  |  |  |  |
| Group | MDD vs *HC* | ↓ (0.83) | *n.s.* | *n.s.* | *n.s.* | ↑ (1.14) |
| Episode status | FE vs *HC* | ↓ (0.78) | *n.s.* | *n.s.* | *n.s.* | ↑ (1.20) |
|  | RE vs *HC* | ↓ (0.84) | *n.s.* | *n.s.* | *n.s.* | ↑ (1.15) |
|  | FE vs *RE* | *n.s.* | *n.s.* | *n.s.* | *n.s.* | - |
| Remission status | ACUTE vs *HC* | ↓ (0.78) | *n.s.* | *n.s.* | *n.s.* | ↑ (1.22) |
|  | REM vs *HC* | *n.s.* | *n.s.* | *n.s.* | *n.s.* | ↑ (1.27) |
|  | ACUTE vs *REM* | *n.s.* | *n.s.* | *n.s.* | ↓ (0.85) | - |
| Antidepressant use | AD use vs *HC* | ↓ (0.88) | *n.s.* | *n.s.* | ↓ (0.87) | ↑ (1.14) |
|  | AD free vs *HC* | ↓ (0.90) | *n.s.* | ↑ (1.03) | *n.s.* | ↑ (1.13) |
|  | AD use vs *AD free* | *n.s.* | *n.s.* | *n.s.* | ↓ (0.88) | *n.s.* |
| Age of onset of MDD | EAO vs *HC* | ↓ (0.78) | *n.s.* | ↑ (1.04) | *n.s.* | ↑ (1.21) |
|  | LAO vs *HC* | ↓ (0.84) | *n.s.* | - | (0.88) | *n.s.* |
|  | LOA vs *EOA* | *n.s.* | *n.s.* | ↓ (0.96) | *n.s.* | ↓ (0.89) |
| Severity | BDI | ↓ (-0.99*pt*) | *n.s.* | *n.s.* | *n.s.* | ↑ (0.82*pt*) |
|  | HDRS | *n.s.* | ↓ (-0.23*pt*) | *n.s.* | ↓ (-0.55*pt*) | *n.s.* |
| BMI* | BMI | ↓ (-0.99*pt*) | ↓ (-0.19*pt*) | ↑ (0.16*pt*) | ↓ (-0.66*pt*) | ↑ (0.48*pt*) |
| Childhood Trauma** | CTQ | *n.s.* | *n.s.* | *n.s.* | *n.s.* | *n.s.* |

### 3.5. Average z-scores

Lower average z-scores were associated with a higher likelihood to be in the MDD group (0.83 relative risk (RR), p<0.001) and being acutely depressed (RR=0.78; p<0.001), compared to being in the HC group. A one unit decrease, equivalent to one standard deviation (SD), in average z-score was associated with a 0.99-point increase in BDI scores (p=0.04), while no association with HDRS scores was observed. A one SD decrease in average z-scores was associated with a 0.99-point BMI (kg/m^2) increase across MDD and HC (p<0.001, see **S1.14**).

### 3.6. Load scores

There were no differences between MDD and HC for positive or negative load scores (**Table 1 & S1.15-16**). For negative load, having more regions with extreme negative deviations was associated with a higher likelihood of not taking antidepressants at time of scanning (vs. HC: RR=1.03; p=0.048) and of having an early onset of MDD (vs HC: RR=1.04, p=0.05, vs. late onset MDD: RR=0.96, p=0.05). A one-unit decrease in positive load scores was associated with a 0.24-point decrease in HDRS scores (p=0.05). Across groups a one unit decrease in positive load scores led to a 0.19 km/m^2 increase in BMI (p=0.01), while a one unit increase in negative load scores was associated with a 0.16 km/m^2 increase in BMI (p = 0.048).

### 2.7. Extremity scores

Positive and negative extremity were defined as the most positive or negative individual z-score across all regions. Greater negative extremity scores were associated with a higher likelihood of being in the MDD group (vs. HC: RR=1.14, p=0.001), of having an early onset of MDD (vs. HC: RR=1.21, p<0.001, vs. late onset: RR=0.89 p=0.02), being either acutely depressed or remitted (acute vs. HC: RR=1.22 p<0.001, remitted vs. HC: RR=1.27 p<0.001), using or not using antidepressants (AD use vs. HC: RR=1.14 p<0.001, AD free vs. HC: RR=1.13 p=0.01) and having higher BDI (0.823-point increases, p=0.04) and BMI scores (0.49-point increase, p=0.04, across groups). Greater positive extremity scores showed a link to a lower likelihood of being in the antidepressant user group (vs. HC: RR=0.87, p=0.009, vs. antidepressant-free group: RR=0.88, p=0.03), of being in the late onset MDD group (vs. HC: RR=0.89, p=0.020), and of being in the acutely depressed group (vs. remitted: RR=0.84, p=0.038). Positive extremity scores were negatively associated with disease severity (-0.56-point HDRS decrease per unit increase) and with BMI (-0.109 unit decrease in BMI, across groups, see **Table 1 & S1.17-18**).

## 4. Discussion

In this large study using NM, we used our previously developed algorithm^43,44^ to characterise the heterogeneity of CT in depression. We found lower z-scores in individuals with MDD compared to HC, mostly in temporal, frontal and parietal areas, and in the insular cortex, with small effect sizes (Cohen’s *d* ranging 0.05-0.2) and with a large overlap in z-scores between MDD patients and HC (distributional overlap ranging 92-98%). The magnitude of these effects and the anatomical distribution of the deviations are in line with our previous results from group average comparison studies^12,14,15^.

### 4.1. Heterogeneity in cortical thickness deviations in depression

The main aim of our NM approach was to examine interindividual differences in CT alterations between individuals with MDD. Assigning either a supra-normal, infra-normal or normal z-score per individual per cortical region revealed that ∼70% of individuals with MDD showed no extreme deviation in any brain region, thus having CT values that fall fully within the norm. The remaining ∼30% of individuals with MDD, however, exhibited wide-spread heterogeneity in degree and spatial distribution of these extreme z-scores. No single cortical region showed an extreme deviation in more than 12.1% of individuals with MDD. This lack of spatial overlap in CT alterations demonstrates the large variation of CT abnormalities in MDD and is in line with spatial percentage overlap in CT z-scores observed in previous NM studies in other disorders such as schizophrenia, ADHD and bipolar disorder^33–35,56^. The variation in extreme deviations in different cortical regions across individuals with MDD may be associated with distinct behavioural and symptom phenotypes that have been reported within MDD^57,58^. Importantly, the high degree of regional heterogeneity within MDD suggests that group average comparisons are not representative of a specific profile of CT alterations apparent in every individual patient (see also case study examples in **S2.8**).

Even though we observed considerable heterogeneity in the spatial location of CT alterations at the regional level they may still form part of the same higher order functional circuit. This is supported by findings showing that regional extreme deviations from NM can be coupled into common higher-order functional circuits and networks^59^. Regional overlap between individuals in extreme z-scores derived from NM increased from <7% up to 21% when integrated into higher order functional networks, in particular in circuits including the prefrontal cortex (PFC) and other frontal areas^59^. This network character of CT alterations could be the driving underlying factor for clinical phenotypes in MDD, re-integrating the heterogeneity at the level of regional CT alterations.

### 4.2. Large overlap in z-score distributions of cortical thickness measures between individuals with depression and healthy controls

Importantly, we found a high similarity between individuals with MDD and HCs in CT measures. The maximum group difference between distributions in z-scores of regional cortical thicknesses was 8%. In other words, 92% of the distributions in CT z-scores overlapped between MDD and HC. Similar findings have been reported previously^11^, with distributional overlap scores of structural brain alterations between healthy individuals and individuals with MDD of between 87% and 95%.

Around 30% of HCs also showed extreme deviations in CT, which was similar to the percentage of individuals with MDD with extreme deviations and reflects the high distributional overlap between MDD and HC. It should be noted, however, that this percentage rises with a larger number of regions analyzed. Given this high overlap between MDD and HC, CT alterations may be driven by other factors commonly associated with MDD but present across individuals with and without depression, such as genetic^60,61^, sociodemographic^62^, clinical (e.g. childhood trauma^63^), nutritional^64^, activity-related^65,66^, and lifestyle factors (e.g., smoking, drinking^67,68^). This hypothesis is supported by our observed negative association between BMI and z-scores of CT alterations across HC and MDD individuals, without any interaction between BMI and diagnosis. A similar association between obesity and CT alterations has been found previously in healthy adults^69–74^. Several studies suggest that increased neuroinflammation may be a mediating factor in the association between obesity and cortical thinning^75–77^ (although also thickening has been reported^76^) and might contribute to the pathophysiology of cortical thickness alterations in depression^70^.

### 4.3. Associations with clinical characteristics within MDD

While one implication of this study is that extreme negative deviations to some extent differentiate between healthy controls and depressed individuals, we also found support that extreme deviations can be used to distinguish between subgroups within depression.

Within depression, having higher positive extremity scores decreased the likelihood of being acutely depressed compared to being remitted. This suggests that slight alterations in CT may normalise following remission, which is in line with previous studies showing attenuated reductions in prefrontal and anterior cingulate cortices and the hippocampus in remitters at a 3-year follow up compared to non-remitters^78^. Alternatively, more extreme positive deviations in CT may protect against chronic depression (or develop in a compensatory manner), as increased orbitofrontal and inferior temporal volumes^79^ and thicker frontal^80^ and anterior cingulate^79^ cortices have been shown to predict future remission.

We further found that, within MDD, more extreme positive CT deviations (greater positive extremity scores) were associated with a decreased likelihood of taking antidepressants. While antidepressants have been associated with neuroprotective effects and neurogenesis^81^ (see also^12,21^), large-scale cross-sectional studies have consistently shown most wide-spread effects with largest effect sizes of structural brain alterations in those with MDD taking antidepressants. This may in part be driven by overall illness severity, as individuals who are more severely depressed or have had a more chronic or recurrent course of depression are more likely to use antidepressants. This is indirectly supported by our finding of negative associations of positive extremity and load scores with overall severity of depressive symptoms (HDRS scores). Nonetheless, longitudinal studies are required to clarify these associations.

Further, more extreme negative deviations were associated with an earlier age of MDD onset (<21 years of age) compared to adult age of onset in MDD. This could either point to more extreme negative deviations in CT constituting a risk factor for the development of depression, or a consequence of longer illness duration (co-correlated with earlier age of onset), which have both been reported previously^82^.

Unlike previous studies, we did not find any association between summary scores of deviations and childhood trauma^63,83^. This discrepancy may stem from the different approach of our study compared to previous studies^63^, as we used summary scores of extreme deviations across cortical regions instead of examining scores per cortical region^83^. In addition, the age-corrected z-score measures used in this study do not allow testing for interactions between CT and age, which have been reported previously^63^, and the inclusion of the total CTQ scores did not allow for any interaction with the subscales of the measurement.

### 4.4. Limitations

Despite being the largest NM study in MDD to date, our study is limited by the cross-sectional nature of the current dataset, not allowing us to determine a causal direction for any of the observed associations. The use of longitudinal data to establish normative trajectories could lead to an increased understanding of the causal processes underlying the association between z-score deviations and clinical variables.

Further, the current study is limited by the clinical covariates currently available in the ENIGMA MDD data set. Detailed information on illness history, genetic and environmental risk factors^61,84^ and duration, type and dosage of antidepressant treatment^15^, were not available for many sites. The inclusion of those parameters into the normative model could be the target of future studies.

## 5. Conclusions and future directions

While most individuals with MDD do not show extreme deviations in CT, the spatial location and distribution of those who do show extreme alterations is highly heterogeneous across individuals. The similarity of the percentage of individuals with extreme deviations in MDD and HC groups points to an important role of non-depression specific mechanisms affecting CT in both MDD and HC (e.g., BMI). Extreme negative deviations predicted MDD group membership, while extreme positive deviations were linked to more favourable clinical characteristics within MDD, though these differences were subtle. This suggests that regional CT may not provide diagnostic biomarkers for MDD. Likely we need more temporally and spatially finely grained imaging techniques to detect structural brain deviations that can better differentiate between MDD and HC or use data-driven techniques that demarcate biotypes. Other neuroimaging modalities, such as functional brain measures, may have the potential to yield larger effect sizes (although see^85^). This limitation in diagnostic sensitivity, however, does not necessarily limit the use of neuroimaging measures as markers of treatment response^85^ or general disease prognosis^86^.

## 6. Author contributions

J.M.M.B: Conceptualization, Writing - Original Draft, Methodology, Software, Formal analysis, Validation, Visualisation, Data curation; L.S.v: Writing - Review & Editing; E.P: Writing - Review & Editing, Project administration, Supervision, Data curation; C.D: Writing - Review & Editing; L.K.M.H: Writing - Review & Editing; S.E.E.C.B: Data curation, Project administration; J.B: Resources, Data curation; F.B: Writing - Review & Editing; K.B: Resources, Data curation, Writing - Review & Editing, Funding acquisition; L.M.B: Writing - Review & Editing;K.B: Data curation, Writing - Review & Editing; R.B: Methodology, Writing - Review & Editing; B.C: Data curation, Writing - Review & Editing; K.R.C: Investigation, Resources, Writing - Review & Editing, Funding acquisition; U.D: Investigation, Resources, Data curation, Writing - Review & Editing, Funding acquisition; D.D: Formal analysis, Resources, Data curation, Writing - Review & Editing, Supervision; K.D: Resources, Data curation; J.W.E: Formal analysis, Data curation C.H.Y.F: Investigation, Resources, Writing - Review & Editing; P.F: Investigation, Data curation, Writing - Review & Editing; B.R.G: Writing - Review & Editing; I.H.G: Writing – Review & Editing, J.G: Investigation, Data curation, Writing - Review & Editing; A.G: Investigation; R.G: Resources, Writing - Review & Editing; H.J.G: Resources, Data curation, Writing - Review & Editing, Project administration; N.A.G: Data curation, Writing - Review & Editing, Project administration; D.G: Data curation, Writing - Review & Editing; O.G: Formal analysis, Investigation, Writing - Review & Editing, Supervision; T.H: Data curation, Writing - Review & Editing; G.B.H: Formal analysis; J.H: Investigation, Resources, Data curation, Writing - Review & Editing; B.J. H: Investigation; S.N.H: Resources, Data curation; M.H: Writing - Review & Editing, Project administration; I.B.H: Methodology, Formal analysis, Data curation, Writing - Review & Editing, Visualization, Funding acquisition; T.C.H: Formal analysis, Data curation, Writing - Review & Editing, Funding acquisition; N.J: Methodology, Resources, Writing - Review & Editing, Project administration; A.J.J: Writing - Review & Editing; A.J: Data curation, Writing - Review & Editing; T.K: Resources, Data curation; T.K: Data curation, Writing - Review & Editing T.T.J.K: Resources, Writing - Review & Editing, Supervision, Project administration, Funding acquisition B.K: Writing - Review & Editing B.K: Writing - Review & Editing A.K: Data curation, Writing - Review & Editing A.K: Data curation, Writing - Review & Editing, Funding acquisition E.J.L: Investigation, Data curation, Writing - Review & Editing R.L: Data curation, Writing - Review & Editing M.L: Data curation, Writing - Review & Editing A.M: Writing - Review & Editing S.E.M: Data curation, Writing - Review & Editing S.M: Data curation, Writing - Review & Editing E.M: Writing - Review & Editing B.M: Visualization I.N: Conceptualization, Resources, Data curation, Project administration, Funding acquisition G.O: Resources, Data curation; M.L.O: Conceptualization, Methodology, Software, Validation, Data curation, Writing - Review & Editing; M.J.P: Writing - Review & Editing, Visualization; E.R: Writing - Review & Editing; L.R: Writing - Review & Editing; P.G.P.R: Writing - Review & Editing; M.D.S: Data curation; R.S: Writing-Review & Editing; P.G.S: Validation, Writing - Review & Editing; H.S: Resources; K.S: Investigation, Resources, Writing - Review & Editing, Funding acquisition; E.S: Writing - Review & Editing; J.C.S: Supervision; F.S: Data curation, Writing - Review & Editing; D.J.S: Writing - Review & Editing; A.S: Writing - Review & Editing; B.S: Resources, Writing - Review & Editing, Funding acquisition; L.T.S: Formal analysis; L.T: Data curation, Writing - Review & Editing; F.T: Data curation, Writing - Review & Editing; S.I.T: Writing - Review & Editing, Project administration; P.U: Data curation, Writing - Review & Editing; N.J.A.v: Resources, Writing - Review & Editing, Funding acquisition; H.V: Investigation, Resources, Data curation, Writing - Review & Editing, Project administration, Funding acquisition; M.W: Resources, Data curation, Funding acquisition; H.C.W: Project administration; S.W: Resources, Writing - Review & Editing, Funding acquisition; N.R.W: Data curation, Writing - Review & Editing; K.W: Data curation; M.W: Data curation; T.T. Y: Resources, Data curation, Writing - Review & Editing, Funding acquisition; C.A.Z: Methodology; G.B.Z: Supervision; P.M.T: Conceptualization, Methodology, Investigation, Resources, Writing - Review & Editing, Supervision, Project administration; D.J.V: Funding acquisition, Data curation; A.F.M: Writing - Review & Editing, Project administration, Supervision, Funding acquisition; L.S: Writing - Review & Editing, Project administration, Supervision, Funding acquisition;

## 7. Acknowledgements

The authors declare the following funding agencies and bodies: ENIGMA MDD is supported by NIH RO1 MH129742, RO1 MH129832 and RO1 MH117601 grants. J.M.M.B: none. L.S v.V: none. E.P: none. C.D: none. L.K.M.H: Nederlandse Organisatie voor Wetenschappelijk Onderzoek (NWO) 09150162210201. S.E.E.C.B: none. J.B: none. F.B: Italian ministry of health RF-2018-12367249. K.B: The BiDirect Study is supported by grants of the German Ministry of Research and Education (BMBF) to the University of Muenster (01ER0816 and 01ER1506). L.M.B: none. K.B: none. R.B: The SHIP study has received funding from the following institutions: Federal Ministry of Education and Research, the Ministry of Cultural Affairs as well as the Social Ministry of the Federal State of Mecklenburg-West Pomerania. Magnetic resonance imaging examinations were supported by Siemens Healthineers, Siemens Healthcare GmbH (Erlangen, Germany). B.C: BCD is funded by the Australian National Health and Medical Research Council CJ Martin fellowship (grant number: 1161356). K.R.C: The study was funded by the National Institute of Mental Health (K23MH090421), the National Alliance for Research on Schizophrenia and Depression, the University of Minnesota Graduate School, the Minnesota Medical Foundation, and the Biotechnology Research Center (P41 RR008079 to the Center for Magnetic Resonance Research), University of Minnesota, and the Deborah E. Powell Center for Women’s Health Seed Grant, University of Minnesota. U.D: This work was funded by the German Research Foundation (grant FOR2107 DA1151/5-1, DA1151/5-2, DA1151/9-1, DA1151/10-1, DA1151/11-1 to UD; SFB/TRR 393, project grant no 521379614) and the Interdisciplinary Center for Clinical Research (IZKF) of the medical faculty of Münster (grant Dan3/022/22 to UD). D.D: none. K.D: none. J.W.E: * Research Support This research was supported (in part) by the Intramural Research Program of the NIMH: ZIAMH002927. C.H.Y.F: Research grant funding on behalf of the University of East London from Flow Neuroscience (R102696); research grant funding: NIMH (R01MH134236), Baszucki Brain Research Fund Milken Institute (BD0000009), Rosetrees Trust (CF20212104), International Psychoanalytic Society (158102845), MRC (G0802594), NARSAD, Wellcome Trust. Associate Editor of Psychoradiology, Section Editor of Brain Research Bulletin. P.F: CIBERSAM, AGAUR, "la Caixa" Foundation (LCF/BQ/PR22/11920017). B.R.G: MRC. J.G: none. A.G: none. R.G: German Federal Ministry of Education and Research (Bundesministerium für Bildung und Forschung, BMBF: 01 ZX 1507, “PreNeSt - e:Med”). H.J.G: none. N.A.G: none. D.G: none. I.H.G: NIH grant R37MH101495. O.G: none. T.H: none. G.B.H: none. J.H: This work was supported by the ALF Grants, Region Östergötland, Sweden. B.J. H: BJH and CGD acknowledge that data collected in Melbourne, Australia, was supported by Australian National Health and Medical Research Council of Australia (NHMRC) Project Grants 1064643 (principal investigator, BJH) and 1024570 (principal investigator, CGD). BJH and CGD were supported by NHMRC Career Development Fellowships (1124472 and 1061757, respectively). S.N.H: none. M.H: none. I.B.H: NHMRC Research Fellowship: Right care, first time: delivering technology-enabled mental health care to young people at scale 2023-2027. T.C.H: National Institute of Mental Health (K01MH117442). N.J: R01MH117601, R01MH134004. A.J: German Research Foundation (DFG): grants JA 1890/7-1, JA 1890/7-2. T.K: none T.T.J.K: This work was funded by the German Research Foundation (DFG grants FOR2107 KI588/14-1, and KI588/14-2, and KI588/20-1, KI588/22-1 to Tilo Kircher, Marburg, Germany). Biosamples and corresponding data were sampled, processed and stored in the Marburg Biobank CBBMR. B.K: none. A.K: none J.L: none. E.J.L: in part funded by the Deutsche Forschungsgemeinschaft (DFG, German Research Foundation) – Project-ID 521379614 – SFB/TRR 393. R.L: none. M.L: none. A.M: none. S.E.M: NHMRC grants APP1172917 and APP1158127. S.M: none E.M: none. B.M: none. I.N: DFG (grants NE2254/1-2, NE2254/2-1, NE2254/3-1, NE2254/4-1) G.O: JP18dm0307002, JP24wm0625204 M.L.O: none. M.J.P: Grant SGR-Cat2021/00832 from the Agencia de Gestió d’Ajuts Universitaris i de Recerca (AGAUR), Generalitat de Catalunya. E.R: none. L.R: none. P.G.P.R: none. M.D.S: none. R.S: none. P.G.S: none. H.S: none. K.S: This study was supported by National Healthcare Group Research Grant, Singapore (SIG/15012) awarded to KS. E.S: none. J.C.S: none. F.S: none. D.J.S: none. A.S: none. B.S: German Research Foundation (STR 1146/18-1, part of FOR2107) L.T.S: none. L.T: none F.T: none S.I.T: SIT is supported in part by NIH grants R01MH123163, R01MH121246, and R01MH116147. Core funding for ENIGMA was provided by the NIH Big Data to Knowledge (BD2K) program under consortium grant U54 EB020403 to PMT. P.U: none. N.J.A.v: none. H.V: SHIP is part of the Community Medicine Research Network of the University Medicine Greifswald, which is supported by the German Federal State of Mecklenburg-West Pomerania. M.W: none. H.C.W: Wellcome Trust. S.W: Brain & Behavior Research Foundation. N.R.W: none. K.W: SHIP is part of the Community Medicine Research net of the University of Greifswald, Germany, which is funded by the Federal Ministry of Education and Research (grants no. 01ZZ9603, 01ZZ0103, and 01ZZ0403), the Ministry of Cultural Affairs and the Social Ministry of the Federal State of Mecklenburg-West Pomerania. MRI scans in SHIP and SHIP-TREND have been supported by a joint grant from Siemens Healthineers, Erlangen, Germany and the Federal State of Mecklenburg-West Pomerania. M.W: none. T.T. Y: This work was supported by the National Center for Complementary and Integrative Health (NCCIH) R21AT009173, R61AT009864, and R33AT009864 to TTY; by the National Center for Advancing Translational Sciences (CTSI), National Institutes of Health, through UCSF-CTSI UL1TR001872 to TTY; by the American Foundation for Suicide Prevention (AFSP) SRG-1-141-18 to TTY; by UCSF Weill Institute for Neurosciences to TTY; by UCSF Research Evaluation and Allocation Committee (REAC) and J. Jacobson Fund to TTY; by the National Institute of Mental Health (NIMH) R01MH085734 and the Brain and Behavior Research Foundation (formerly NARSAD) to TTY. C.A.Z: National Institute of Mental Health. G.B.Z: none. P.M.T: NIH grant R01 MH131806. D.J.V: none. A.F.M: EC | ERC | HORIZON EUROPE European Research Council (ERC) MENTALPRECISION ”10100118, HHS | NIH | National Institute of Mental Health (NIMH) 1R01MH130362-01A1. L.S: NHMRC Investigator Grant (2017962) and University of Melbourne Dame Kate Campbell fellowship.

