## Supplement S2 for "Dissecting heterogeneity in cortical thickness abnormalities in major depressive disorder: a large-scale ENIGMA MDD normative modelling study"

**SUPPLEMENTARY MATERIAL 2**

**Supplementary Text S2.1 Data cleaning**

**Supplementary Text S2.2 Processing pipeline**

**Supplementary Text S2.2a Evaluation of the created data sets**

**Supplementary Table S2.2a Sex distribution in the training control, test control and test MDD set.**

**Supplementary Table S2.2b Age distribution in the healthy training control, healthy test controls and test MDD set**

**Supplementary Figure S2.3 Sensitivity analysis**

**Supplementary Text S2.4 Modelling**

**Supplementary Text S2.5 Model fit**

**Supplementary Text S2.5a. Analysis of distribution of extreme deviations across sets**

**Supplementary Text S2.6 Alignment of our results with Cohen's d values from previous ENIGMA MDD publication on cortical thickness differences in MDD[3]**

**Supplementary Text S2.7 Calculation of distribution overlap scores.**

**Supplementary Text S2.8 Calculation of percentage overlap scores.**

**Supplementary Figure S2.9 Correlation between summary scores**

**Supplementary Figure S2.10. Profiles of individuals with MDD with the most negative load and extremity score and the most negative average z-score.**

**Supplementary Text 2.11 Expected number of extreme regions per individual and false discovery rate correction for z-scores**

### Supplement S2

#### S2.1 Data cleaning

Region-wise averages between the right and the left hemisphere were created for all 10,454 individuals of the data set. For participants with one missing values in one hemisphere, the final value was created by taking the value from the other hemisphere. This left us with 35 measures of interest: 34 cortical regions, and one average between the entire left and right hemisphere. Out of 10,454 individuals in the ENIGMA MDD data set, a total of 1,509 individuals were excluded: 306 individuals did not have information about disease status and, 440 subjects were excluded as they exhibited more than four missing values across the 35 regions of interest, and 763 individuals were excluded because either i) their corresponding sites did not give consent to be included in the study, ii) their site only contained depressed OR healthy individuals, iii) the entire site was considered an outlier by visual inspection in terms of cortical thickness measures. This left us with 8,945 individuals that were included into the study out of which N=5,300 were healthy controls and N=3,645 were diagnosed with major depressive disorder. The total sample included individuals from 34 sites including 40 different scanners.

#### S2.2 Processing pipeline

The total of N=5,300 healthy controls were divided into a HC training set (N=3,181) and a HC test set (N=2,119) using the R Package caret (https://cran.r-project.org/web/packages/caret/) implementing a 60/40 within-site split. Data from all individuals with MDD that remained after data cleaning were included in into a third set, the MDD test set. Comparing the MDD test set against a healthy control test set instead of against forward predictions from the HC training set allows direct comparison between the sets, as both the healthy control test set and the MDD test set yield the same generalisation error in their predictions (avoiding overfitting). In the created sets, each scanner was represented by at least 5 individuals in the training and test sets. Data imputation, scaling and centering across sites was performed using the *knnImpute* function in *caret* with a radius of k=10 to impute missing data in the MDD test set and the healthy control test set based on the healthy control training set. Consecutively, the normative model was trained on the healthy control training set and subsequently applied to the healthy control and MDD test sets to obtain predicted z-score CT values. Similarly to the imputation procedure, this approach allows a comparison between the healthy control test set and the MDD test set. An overview over the processing pipeline can be found in **Figure 1** of the main text.

#### S2.2a Evaluation of the created data sets

The male/female and age distribution in the HC train, HC test and MDD test sets were compared using a multinomial logistic regression (multinom) from the nnet package in R, in which set (HC train, HC test, MDD test) was predicted by age and sex. The results reflect the choices made to generate those sets: the baseline odds to be in the HC test set were 37% lower than being in the HC train set ($\beta = -0.49$, with odds test controls = exp($\beta$) = 0.61, p < 0.001) which reflects the 40/60 split of generating the healthy control training and test sets. Further, the and increased percentage of female individuals with depression was reflected in increased effect of age on odds of being the MDD test set, which was stronger for women than for men (significant Age:sex interaction, $\beta$ = 0.01,p < 0.001). There was no significant difference in the any other odds of age or sex of predicting the HC test versus MDD test set. See an overview of the sex and age distribution in table **S2.2a and S2.2b**)

| | **Training set healthy controls** | **Test set healthy controls** | **Test set MDD** |
|---|---|---|---|
| Male | 1386 (44%) | 923 (44%) | 1344 (37%) |
| Female | 1795 (56%) | 1196 (56%) | 2301 (63%) |
| Total | 3181 (100%) | 2119 (100%) | 3645 (100%) |

**Table S2.2a.** Sex distribution in the training control, test control and test MDD set. Percentages of male/female distribution in brackets.

| | [0,10) | [10,20) | [20,30) | [30,40) | [40,50) | [50,60) | [60,70) | [70,80] | [80,90) |
|---|---|---|---|---|---|---|---|---|---|
| Training set, healthy controls | 20 (72%) | 292 (31%) | 1015 (43%) | 377 (33%) | 538 (31%) | 523 (32%) | 308 (38%) | 99 (42%) | 9 (36%) |
| Test set, healthy controls | 4 (14%) | 182 (20%) | 633 (27%) | 273 (24%) | 382 (22%) | 358 (22%) | 206 (26%) | 75 (32%) | 6 (24%) |

| | | | | | | | | | |
|---|---|---|---|---|---|---|---|---|---|
| Test set, MDD | 4 (14%) | 462 (49%) | 728 (31%) | 485 (42%) | 831 (47%) | 773 (47%) | 291 (36%) | 61 (26%) | 10 (40%) |
| Total | 28 (100%) | 936 (100%) | 2376 (100%) | 1135 (100%) | 1751 (100%) | 1025 (100%) | 805 (100%) | 235 (100%) | 25 (100%) |

**Table S2.2b:** Age distribution in the healthy training control, healthy test controls and test MDD set. Percentages with reference to the age bin in brackets

**S2.3 Sensitivity analysis**

In order to estimate the effect of imputing the MDD test set by the healthy control training set, we further performed a sensitivity analysis by imputing the MDD test set on itself (k=10 nearest neighbour imputation). We report differences in the means of the MDD test sets (imputed by the healthy control training set or by itself) for 13 out of 35 regions (see for a detailed overview and mean difference comparison **S1.19**).

**S2.4 Modelling**

A hierarchical Bayesian regression model with age, sex and site as predictors was used to create a model mapping the normative variation in each CT measure onto age and sex, while accommodating for multi-site effects. As the inversion of the Gaussian Process matrix is computationally costly and the computation time scales quadratically with N (O($n^2$)), we used a B-splines model with 5 knots to model non-linear associations between age and CT.

The predictions from this model allowed us to calculate z-scores for each participant *i* and region *j* using the following:

$$z_{ij} = \frac{f(x_{ij}) - y_{ij}}{\sqrt{\sigma^2 + \sigma_*^2}}$$

Where $f(x_{ij})$ is the prediction of the model, $y_{ij}$ is the true value, $\sigma^2$ is a random gaussian noise component and $\sigma_*^2$ is the predictive variance from the model (for more details see [1,2]).

**S2.5 Model fit**

We calculated standardized the Pearson correlation coefficient ρ root mean square errors (SRMSE), the Expected Variation (EV)and the mean standardized log loss (MSLL) for each region for the HC training set, the HC test set and the MDD test set. The mean SRMSE for the HC training set was 0.69, for the HC test set 0.70 and for the MDD test set 0.71, which demonstrates good fits. Similarly, the correlation coefficient ρ was satisfactory for all sets (mean EV HC training set = 0.50; mean EV HC test set = 0.49; mean EV MDD test set = 0.41) as well as the explained variance (mean ρ HC training set = 0.71; mean ρ HC test set = 0.69; mean ρ MDD test set = 0.64). The closeness for those values between training and MDD and control test set also demonstrate a good generalisation of the training models to both test sets. In addition, the MSLL shows that predictions of the model for the HC test set and MDD test set were better than predictions made from the mean of the training set (MSLL: MDD: -0.18. Test HC: -0.25). A complete breakdown of all model fit parameters per region can be found in the supplementary tables **S1.10-S1.13.**

**S2.5a. Analysis of distribution of extreme deviations across sets.**

Further assessing the model fit, we examined the distribution of predictive z-scores against the Gaussian distribution it should be following. Hence the percentage of extreme deviations in the HC training set should be around 2.5% for $z < -1.96$ and $z > 1.96$. We have extracted the distributions of extreme deviations for the HC training set, the HC test set and the MDD test set in table S1.9.

### S2.6 Alignment of our results with Cohen's d values from previous ENIGMA MDD publication on cortical thickness differences in MDD[3]

We further evaluated the alignment of the group differences extracted from the z-scores in our analysis (Cohen's d estimates) with Cohen's d group differences from our previous publication[3]. As Cohen's d estimates from the previous publication had been calculated for each hemisphere separately, we first created one average effect size per region across the two hemispheres. Spearman rank & Pearson correlation tests using the *cor.test* function as implemented in the *stats* package in R was used to determine the correlation between ours and previous results. The results showed substantial alignment (Spearman correlation: $\rho$ = 0.67, S = 2362.3, $p < 0.0001$; Pearson correlation: $\rho$ = 0.70, t = 5.6594, df = 33, p-value < 0.0001). The association is also plotted in Figure S2.6 below.

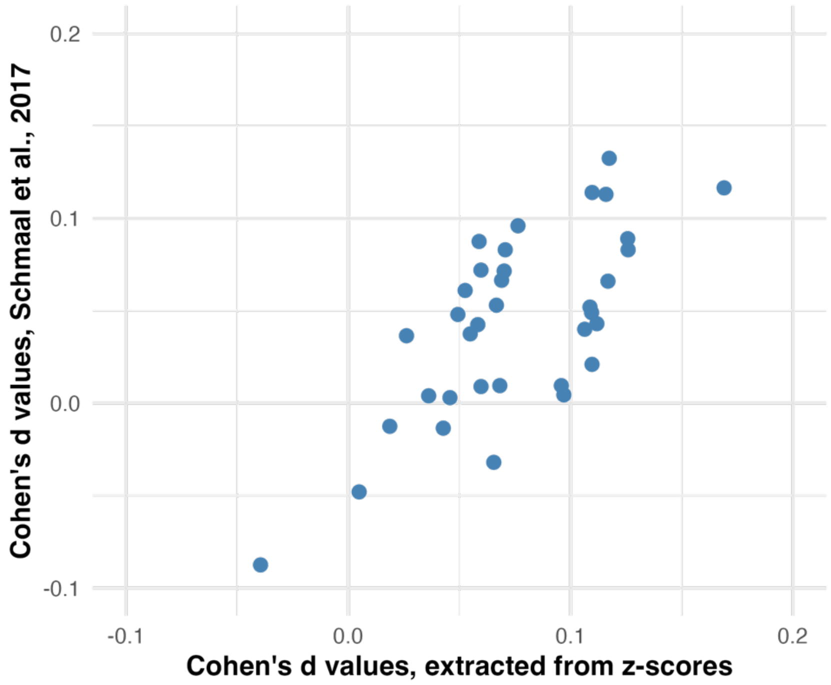


Figure S2.6. Correlation between Cohen's d values extracted from z-scores (current analysis) and Cohen's d scores from previous publication [3].

#### S.2.7 Calculation of distribution overlap scores.

Distributional overlap scores were calculated based on a previous publication[4]. In brief, Gaussian distributions (mean, standard deviations) were fit to the z-score distributions of healthy controls and individuals with depression per region, allowing to calculate the distribution overlap per region. More details can be found in[4].

#### S.2.8 Calculation of percentage overlap scores.

Percentage overlap scores are calculated as a measure of the distribution of extreme deviations across the brain, separately for positive and negative extreme deviations and separately for healthy controls and individuals with depression. The percentage overlap score for a specific region is hence calculated as a fraction between the number of extreme deviations in that region and the number of individuals with at least one extreme deviation:

$$\frac{n_{extreme\_deviations\_region}}{n_{individuals\ with\ at\ least\ one\ extreme\ deviation}}$$

An overview of percentage overlaps for each region, for extreme positive and negative deviations and healthy controls and individuals with depression is given in table **S1.8**. Note that since an individual can show extreme deviations in multiple regions the percentages do not add up to 100%.

#### S2.9 Correlation between summary scores

The analysis of the correlation between summary scores revealed that they were all positively and significantly correlated (all $p < 0.0001$). See table **S2.7**.

| Positive extremity | Negative load | Negative extremity | Average z-score | |
|---|---|---|---|---|
| 0.644 | 0.115 | 0.322 | 0.594 | Positive load |
| | 0.320 | 0.447 | 0.763 | Positive extremity |
| | | 0.671 | 0.619 | Negative load |
| | | | 0.620 | Negative extremity |

**Table S2.9:** Correlation coefficients between averaged z-score, load scores and extremity scores. The p-values of all correlations were statistically significant ($p < 0.05$).

#### S2.10 Profiles of individuals with MDD with the most negative load and extremity score and the most negative average z-score

A

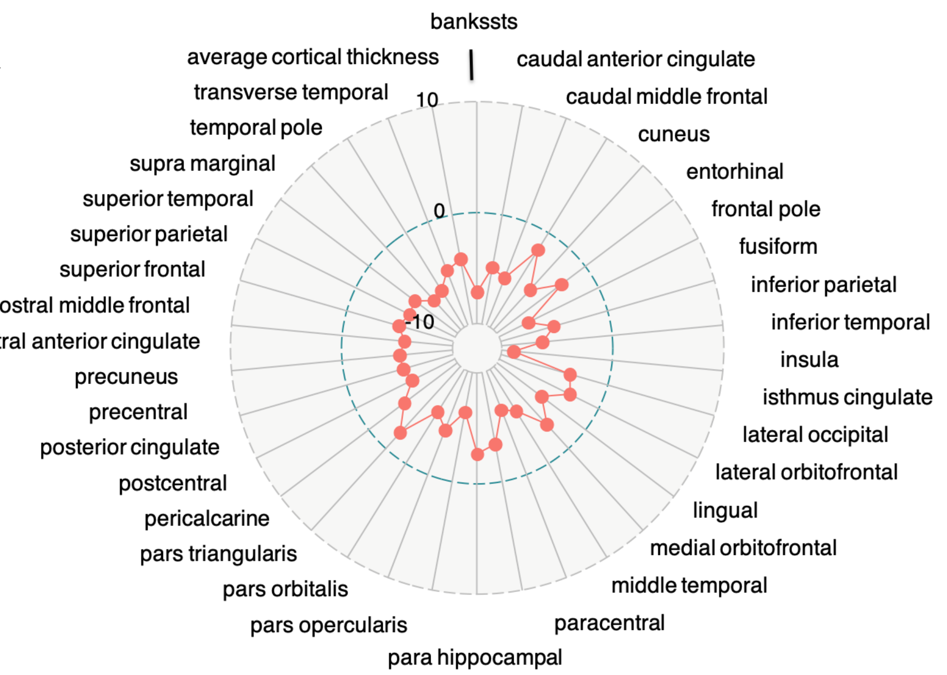


| Subject 3002 | |
|---|---|
| Sex | Male |
| Age | 80 |
| AO | 79 |
| Recur | 2 |
| AD | 1 |
| Rem | NA |
| HDRS | NA |
| BMI | 25 |
| BDI | 22 |
| CTQ | 31 |
| ExN | 33 |
| ExP | 0 |

B

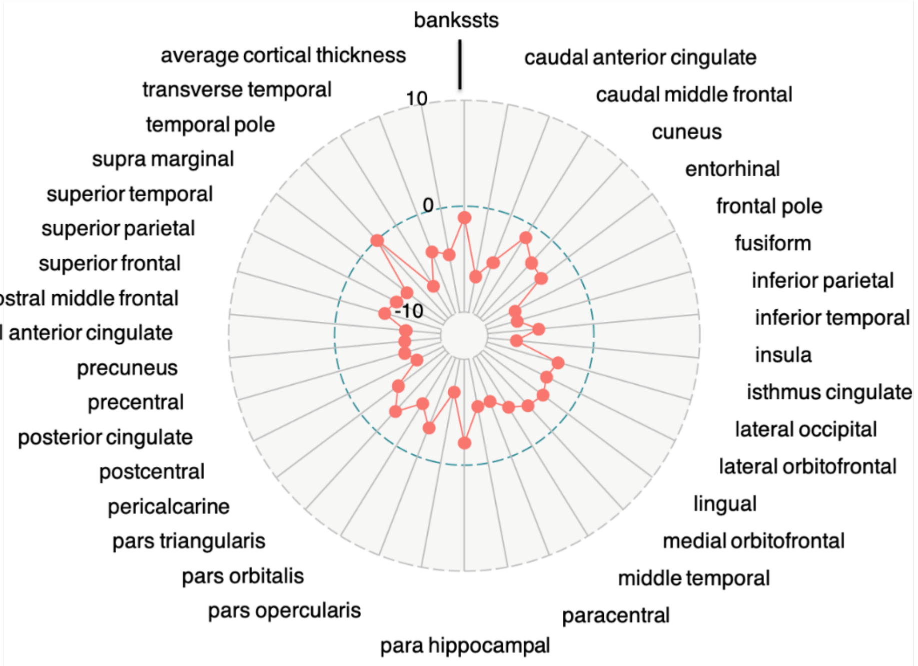


| Subject 3473 | |
|---|---|
| Sex | F |
| Age | 78 |
| AO | 50 |
| Recur | 2 |
| AD | 2 |
| Rem | 1 |
| HDRS | 0 |
| BMI | NA |
| BDI | NA |
| CTQ | NA |
| ExN | 32 |
| ExP | 0 |

**Figure S2.8.** Individualized profiles of individuals with MDD. A) the person with both the most negative load (-33) and the most negative extremity score (-8.927). B) the person with the smallest (most negative) average z-score

Legend: AO: Age of onset of depression. Recur: Number of recurrent episodes. AD: Anti-depressant use. Rem: Remission status. HDRS: Hamilton Depression Rating Scale. BMI: Body Mass Index. BDI: Beck Depression Inventory. CTQ: Childhood Trauma Questionnaire. ExN: number of extreme negative deviations. ExP: number of extreme positive deviations.

### S2.11 Expected number of extreme regions per individual and false discovery rate correction for z-scores

Assuming a threshold $\boldsymbol{\alpha}$ defining an extreme deviation, we can calculate the number of regions expected to show an extreme deviation per individual as: $\boldsymbol{\alpha}$ * N_total_regions. In the case presented here, we can expect 0.05 * 35 = 1.75 regions to show an extreme deviation at random. One way to correct for this effect of multiple comparisons would be to use false discovery correction within each individual and adjust $\boldsymbol{\alpha}$ to obtain an individualized $\boldsymbol{\alpha_i}$ per individual. While this approach would correct for multiple comparisons, it has the undesirable effect that the definition of extreme deviation differs between subjects. Further, pervious research showed a minimal difference between corrected and uncorrected z-scores for the comparison of 35 Desikan-Kiliani[5] regions. Our decision to use uncorrected z-scores in downstream analyses is based on these two factors.


1. Bayer, J. M. M. *et al.* Accommodating site variation in neuroimaging data using normative and hierarchical Bayesian models. *Neuroimage* **264**, 119699 (2022).
2. Rasmussen, C. E. & Williams, C. K. I. *Gaussian Processes for Machine Learning*. (MIT Press, 2005).
3. Schmaal, L. *et al.* Cortical abnormalities in adults and adolescents with major depression based on brain scans from 20 cohorts worldwide in the ENIGMA Major Depressive Disorder Working Group. *Mol. Psychiatry* **22**, 900–909 (2017).

4. Winter, N. R. *et al.* Quantifying Deviations of Brain Structure and Function in Major Depressive Disorder Across Neuroimaging Modalities. *JAMA Psychiatry* **79**, 879–888 (2022).

5. Desikan, R. S. *et al.* An automated labeling system for subdividing the human cerebral cortex on MRI scans into gyral based regions of interest. *Neuroimage* **31**, 968–980 (2006).
