## Supplement S1 for "Dissecting heterogeneity in cortical thickness abnormalities in major depressive disorder: a large-scale ENIGMA MDD normative modelling study"

### SUPPLEMENTARY MATERIAL 1

**Supplementary Table S1.1: ENIGMA - Major Depressive Disorder Working Group Demographics. Age (in years), sex, and MDD patients-control breakdown for participating sites and training and test sets.**

**Supplementary Table S1.2: ENIGMA - Major Depressive Disorder Working Group diagnostic measurement and inclusion and exclusion criteria.**

**Supplementary Table S1.3: ENIGMA - Major Depressive Disorder Working Group image acquisition criteria, breakdown by site**

**Supplementary Table S1.4: ENIGMA - Major Depressive Disorder Working Group Clinical characteristics of MDD patients. Mean age of onset, BDI and HDRS values.**

**Supplementary Table S1.5: Percentage of MDD patients using antidepressant medication, percentage of first episode and recurrent episode MDD patients, percentage of acutely depressed and remitted MDD patients, break down per site.**

**Supplementary Table S1.6: ENIGMA - Major Depressive Disorder Working Group Clinical characteristics that were shared across individuals with MDD and healthy controls. BMI (Body Mass Index), CTQ (Children Trauma Score) for individuals with depression and healthy controls breakdown for available participating sites.**

**Supplementary Table S1.7: Z-score group differences between HC and individuals with MDD. Cohen's d; p-values of the difference (FDR corrected); distributional overlap score (range: 0: no overlap; 1: complete overlap), breakdown per region.**

**Supplementary Table S1.8 Percentage overlap: Percentage overlap per region.**

**Supplementary Table S1.9: Percentages of train healthy controls, test healthy controls and individuals with depression with an extreme deviation (positive deviation: z > 1.96; negative deviation z< -1.96), breakdown region-wise**

**Supplementary Table S1.10: Model fit in Stan: Correlation coefficient /rho per set, region-wise breakdown.**

**Supplementary Table S1.11: Model fit in Stan: Standardized root mean squared errors (SRMSE) per set, region-wise breakdown.**

**Supplementary Table S1.12: Model fit in Stan: Explained variance, per set, region-wise breakdown.**

**Supplementary Table S1.13: Model fit in Stan: Mean standardized log loss per set, region-wise breakdown**

**Supplementary Table S1.14: Average z-score predicting clinical variables.**

**Supplementary Table S1.15: Positive load predicting clinical variables.**

**Supplementary Table S1.16: Negative load predicting clinical variables.**

**Supplementary Table S1.17: Positive extremity predicting clinical variables.**

**Supplementary Table S1.18: Negative extremity predicting clinical variables.**

**Supplementary Table S1.19: Comparison of impact of imputation of test MDD set by train healthy controls vs, by test MDD set.**

**S1.1. ENIGMA - Major Depressive Disorder Working Group Demographics. Age (in years), sex, and MDD patients-control breakdown for participating sites and training and test sets.**

| | Train controls | | | | | Test controls | | | | | MDD | | | | | |
|---|---|---|---|---|---|---|---|---|---|---|---|---|---|---|---|---|
| **Site** | mean age (SD) | | n fem | % fem | total | mean age (SD) | | n fem | % fem | total | mean age (SD) | | n fem | % fem | total MDD | n total |
| **ClinG** | 25.1 | 5.1 | 110 | 56 | 196 | 25.1 | 5.4 | 81 | 65 | 125 | 36.3 | 11.6 | 26 | 53 | 49 | 370 |
| **Edinburgh (Bipolar Family Study**) | 23.5 | 2.5 | 24 | 62 | 39 | 22.6 | 2.2 | 14 | 67 | 21 | 23 | 3 | 11 | 61 | 18 | 78 |
| **McMaster University Mood Disorders** | 30.4 | 11.2 | 22 | 63 | 35 | 30.3 | 12.3 | 8 | 62 | 13 | 35.3 | 13.4 | 27 | 53 | 51 | 99 |
| **Houston** | 38.4 | 12.5 | 39 | 68 | 57 | 39.4 | 12.2 | 29 | 67 | 43 | 39.4 | 13.4 | 54 | 71 | 76 | 176 |
| **Münster Neuroimaging Cohort** | 35 | 11.8 | 237 | 61 | 391 | 36.1 | 12.5 | 147 | 57 | 256 | 38.2 | 12 | 145 | 63 | 232 | 879 |
| **Melbourne** | 19.9 | 2.7 | 32 | 60 | 53 | 19.3 | 3.2 | 22 | 48 | 46 | 19.3 | 2.8 | 76 | 54 | 142 | 241 |
| **Stanford T1w Aggregate** | 39 | 10.7 | 17 | 63 | 27 | 36.6 | 11 | 9 | 50 | 18 | 37.4 | 10.5 | 29 | 60 | 48 | 93 |
| **Magdeburg** - Sexpect | 30 | 0.8 | 1 | 25 | 4 | 28.3 | 4.7 | 0 | 0 | 3 | 32 | 7.5 | 1 | 20 | 5 | 12 |
| **BRCDECC London** | 52.4 | 8.4 | 21 | 58 | 36 | 50.7 | 7.2 | 11 | 44 | 25 | 47.9 | 8.9 | 47 | 68 | 69 | 130 |
| **Barcelona** | 46.2 | 7.1 | 16 | 80 | 20 | 45.8 | 10 | 7 | 58 | 12 | 47 | 7.7 | 49 | 79 | 62 | 94 |
| **Houston adolescents** | 12.4 | 2.8 | 27 | 46 | 59 | 13.5 | 2.7 | 10 | 37 | 27 | 12.9 | 2.5 | 10 | 36 | 28 | 114 |
| **EPISCA (Leiden)** | 14.8 | 1.8 | 17 | 94 | 18 | 14.6 | 1.1 | 9 | 75 | 12 | 15.4 | 1.5 | 16 | 84 | 19 | 49 |
| **UCSF** | 15.5 | 1.3 | 30 | 56 | 54 | 15 | 1.2 | 12 | 36 | 33 | 15.6 | 1.4 | 48 | 65 | 74 | 161 |
| **Sao Paulo (Wellcome)** | 30.1 | 8.2 | 26 | 48 | 54 | 30.4 | 8.1 | 14 | 41 | 34 | 29 | 8.3 | 17 | 71 | 24 | 112 |
| **Minnesota** | 15.9 | 2 | 12 | 60 | 20 | 15.5 | 2 | 14 | 70 | 20 | 15.4 | 1.8 | 53 | 76 | 70 | 110 |
| **Calgary** | 15.6 | 5.1 | 23 | 64 | 36 | 16.15 | 4.9 | 6 | 38 | 16 | 16.9 | 2.2 | 31 | 29 | 55 | 107 |
| **QTIM** | 22.1 | 3.0 | 115 | 65 | 177 | 22.1 | 3.0 | 73 | 70 | 105 | 22.1 | 2.7 | 77 | 75 | 102 | 384 |
| **Oxford** | 32.5 | 13.6 | 7 | 54 | 13 | 28.7 | 6.3 | 11 | 61 | 18 | 30.1 | 10.6 | 24 | 63 | 38 | 69 |
| **FOR2107** | 32.74 | 11.88 | 219 | 62 | 356 | 32.67 | 12.1 | 171 | 70 | 244 | 37.6 | 13.5 | 305 | 65 | 470 | 1070 |
| **AFFDIS** | 31.1 | 13.3 | 7 | 54 | 13 | 34.5 | 10.5 | 2 | 33 | 6 | 39.1 | 15.3 | 12 | 43 | 28 | 47 |
| **Singapore** | 39.1 | 5.3 | 6 | 55 | 11 | 37.8 | 3.5 | 2 | 40 | 5 | 40.1 | 7.6 | 10 | 45 | 22 | 38 |
| **ETPB** | 34.1 | 10.7 | 8 | 57 | 14 | 33.6 | 10.5 | 8 | 67 | 12 | 35.9 | 9.6 | 20 | 59 | 34 | 60 |
| **BiDirect** | 52.1 | 8.1 | 122 | 48 | 255 | 52.3 | 8.1 | 94 | 53 | 176 | 48.8 | 7.2 | 341 | 60 | 572 | 1003 |
| **Sydney** | 46.9 | 23.2 | 38 | 58 | 66 | 46.6 | 23.5 | 22 | 56 | 39 | 35.6 | 21.9 | 140 | 66 | 212 | 317 |
| **Moral Dilemma** | 18.8 | 1.8 | 35 | 1 | 35 | 17.6 | 1.4 | 11 | 1 | 11 | 19.4 | 2.2 | 24 | 1 | 24 | 70 |
| **Stanford FAA** | 31 | 11.4 | 14 | 1 | 14 | 28.6 | 5.1 | 4 | 1 | 4 | 35.6 | 8.4 | 14 | 1 | 14 | 32 |
| **FIDMAG** | 43.5 | 12.8 | 14 | 67 | 21 | 49.9 | 8.5 | 8 | 62 | 13 | 49.3 | 12.2 | 22 | 65 | 34 | 68 |
| **SoCAT** | 33.3 | 12.9 | 48 | 87 | 55 | 40.3 | 13.6 | 42 | 93 | 45 | 39.7 | 12.9 | 71 | 90 | 79 | 179 |
| **SHIP-TREND-0** | 50.4 | 14.4 | 240 | 43 | 555 | 51 | 14.1 | 168 | 45 | 373 | 49.2 | 12.2 | 203 | 65 | 312 | 1240 |
| **Hiroshima** | 36.94 | 11.89 | 59 | 63 | 93 | 38.1 | 10.6 | 45 | 59 | 76 | 43.7 | 11.9 | 71 | 47 | 150 | 319 |
| **CSAN** | 33.3 | 13.6 | 20 | 67 | 30 | 33.1 | 9.9 | 14 | 74 | 19 | 35.9 | 13.5 | 40 | 67 | 60 | 109 |
| **SHIP** | 55.7 | 12.5 | 114 | 45 | 255 | 54.6 | 13 | 79 | 45 | 176 | 53.8 | 11.7 | 96 | 71 | 135 | 566 |
| **MPIP** | 49.2 | 12.3 | 75 | 64 | 118 | 50.0 | 12.9 | 49 | 53 | 93 | 48.01 | 13.9 | 191 | 57 | 337 | 548 |

Fem: female

### S1.2: ENIGMA - Major Depressive Disorder Working Group diagnostic measurement and inclusion and exclusion criteria.

| Cohort | Diagnosis measurement | Exclusion criteria |
|---|---|---|
| **ClinG** | ICD-10 interview | MDD subjects: past or actual presence of other axis I diagnoses other than anxiety disorders, alcohol/cannabis abuse and tobacco dependence; neurological or other medical conditions that could be related to affective symptoms<br>Control subjects: no medical history, including neurological and psychiatric history, as well as no previous or actual use of psychotropic medication |
| **Edinburgh (Bipolar Family Study**) | SCID interview | MDD subjects: presence of any other axis I diagnoses.<br>Control subjects: no medical history, including neurological and psychiatric history, as well as no previous or actual use of psychotropic medication All subjects: any major neurological disorder, learning disability, or any history of head injury that included loss of consciousness and any contraindications to MRI. |
| **McMaster University Mood Disorders** | SCID interview | MDD subjects: Comorbid Axis 1 disorders excluded, including for example, psychosis, bipolar, PTSD substance dependence or current active eating disorder. Exclusion criteria included: i) treatment with anti-cholinergic or typical (first generation) anti-psychotic medication; ii) electroconvulsive therapy (ECT) or transcranial magnetic stimulation (TMS) within the past year; iii) a history of substance dependence or significant and recent (< 1 year) substance abuse; iv) a history (within the past 12 months) of an endocrine or other medical disorder known to adversely affect cognition (e.g., Cushing's, uncontrolled diabetes, seizure disorder); and v) English comprehension lower than a grade 6 reading level. |
| **Houston** | SCID interview | MDD subjects: age below 18; lifetime or current diagnosis of psychotic disorder, or bipolar I or II disorder; substance abuse/dependence in 6 months prior to study inclusion; current major medical problems.<br>Control subjects: age below 18; current major medical problems; current psychiatric or neurologic disorder; history of psychiatric disorders in a first-degree relative; current major medical problems. Both groups: MRI contraindications |
| **Münster Neuroimaging Cohort** | SCID interview | MDD subjects: presence of bipolar disorder, schizoaffective disorders and schizophrenia; substance-related disorders or current benzodiazepine treatment (wash out of at least three half-lives before study participation), and former electroconvulsive therapy.<br>Control subjects: any current or former psychiatric disorder. Both groups: any neurological abnormalities, MRI contraindications |
| **Melbourne** | SCID interview | MDD subjects: lifetime or current SCID-I diagnosis of psychotic disorder, or bipolar I or II disorder. Control subjects: any SCID-I diagnosis or medication use. Both groups: Acute or unstable medical disorder; general MRI contraindications |
| **Stanford T1w Aggregate** | SCID interview | MDD subjects: presence of axis-I disorders other than MDD, anxiety and eating disorders.<br>Control subjects: control individuals did not meet diagnostic criteria for any current psychiatric. Both groups: alcohol / substance abuse or dependence within six months prior to MRI scanning, history of head trauma with loss of consciousness > 5 min, aneurysm, or any neurological or metabolic disorders that require ongoing medication or that may affect the central nervous system (including thyroid disease, diabetes, epilepsy or other seizures, or multiple sclerosis), MRI contraindications, or bad MRI data (e.g., extreme movement). |
| **Magdeburg** - Sexpect | ICD-10 interview | MDD subjects: history of seizures, medication with glutamate modulating drugs (ketamine, riluzole, etc.) or benzodiazepines, prior electroconvulsive therapy (ECT) treatments and pregnancy, atypical forms of depression, any additional psychiatric disorder, and a history of substance abuse or dependence.<br>Control subjects**:** psychiatric illness. Both groups: contraindications against MRI, major medical and neurological illness. |
| **BRCDECC London** | SCAN interview | Contraindications to MRI, diagnosis of neurological disorder, head injury leading to loss of consciousness or conditions known to affect brain structure or function (including alcohol or substance misuse), if they or a first-degree relative had ever fulfilled criteria for mania, hypomania, schizophrenia or mood-incongruent psychosis. |

| | | |
|---|---|---|
| **Barcelona** | DSM-IV-TR acc. to CIDI-interview and HAMD | MDD subjects: Axis I comorbidity according to DSM-IV-TR criteria<br>Control subjects: lifetime psychiatric diagnoses, first-degree relatives with psychiatric diagnoses and clinically significant physical or neurological illnesses, Axis I comorbidity according to DSM-IV-TR criteria |
| **Houston adolescents** | Major depressive disorder (MDD) diagnosis according to DSM-IV | MDD subjects: head trauma with residual effects, neurological disorders, uncontrolled major medical conditions based on patient self-reports and current drug abuse. In addition,<br>Control subjects: history of any Axis I disorder or had a first-degree relative with any Axis I disorder. |
| **EPISCA (Leiden)** | ADIS | MDD subjects: Primary DSM-IV clinical diagnosis of ADHD, ODD, CD, pervasive developmental disorders, post-traumatic stress disorder, Tourette's syndrome, obsessive–compulsive disorder, bipolar disorder, and psychotic disorders; current substance abuse; history of neurological disorders or severe head injury; age < 12 or > 21 years; pregnancy; left-handedness; IQ score < 80 as measured by the Wechsler Intelligence Scale for Children (WISC) (Wechsler, 1991) or Adults (Wechsler, 1997); and general MRI contraindications. |
| **UCSF** | KSADS (semi-structured interview based on DSM) for MDD, DISC/DPS for HCL | All subjects: 1) use of pharmacotherapeutics for treating psychiatric conditions within the past 6 months, 2) misuse of drugs within two months prior to MRI scanning; 3) two or more alcoholic drinks per week within the previous month (as assessed by the Customary Drinking and Drug Use Record; CDDR) (Brown et al, 1998); 4) a full scale IQ score of less than 75 (as assessed by the Wechsler Abbreviated Scale of Intelligence; WASI) (Wechsler, 1999); 5) contraindications for MRI including ferromagnetic implants and claustrophobia; 6) pregnancy or the possibility of pregnancy; 7) left-handedness; 8) prepubertal status (as assessed as Tanner stages of 1 or 2) (Tanner, 1962); 9) inability to understand and comply with procedures; 10) neurological disorder (including meningitis, migraine, or HIV); 11) head trauma; 12) learning disability; 13) serious health problems; and 14) complicated or premature birth (i.e., birth before 33 weeks of gestation).<br>MDD subjects: primary psychiatric diagnosis other than MDD.<br>Control subjects 1) history of mood or psychotic disorders in a first- or second-degree relative (as assessed by the Family Interview for Genetics; FIGS) (Maxwell, 1992); and 2) current or lifetime DSM-IV-TR Axis I psychiatric disorder. |
| **Sao Paulo (Wellcome)** | Hamilton Rating Scale for Depression (HRSD) | MDD subjects: psychotic disorders due to a general medical condition or substance-induced psychosis were excluded. Additional exclusion criteria were: (a) history of head injury; (b) presence of neurological disorders or any organic disorders that could affect the central nervous system; and (c) contraindications for MRI.<br>Control subjects: personal history of psychosis or other Axis I disorders, except substance misuse or mild anxiety disorders. |
| **Minnesota** | SADS; (CDRS-R). | MDD subjects, control subjects: Exclusion criteria for both groups included the presence of a neurologic or other chronic medical condition, mental retardation, pervasive developmental disorder, substance use disorder, bipolar disorder, or schizophrenia |
| **Calgary** | KSADS | MDD subjects: A history of neurological illness, medical illness, claustrophobia, >21 year of age, or the presence of a ferrous implant or pacemaker. Control subjects: Left handed; history of seizures, epilepsy or other neurological or psychiatric diagnoses (specifically bipolar disorder, psychosis, pervasive developmental disorder, eating disorders, PTSD); pregnancy |
| **QTIM** | CIDI interview | MDD subjects: presence of axis-I disorders other than MDD and anxiety disorders<br>Control subjects: antidepressant use, psychiatric disorders All subjects: relatedness between subjects, left handedness, history of neurological or other severe medical illness, head injury or current or past diagnosis of substance abuse, use of cognition affecting medication and general MRI contraindications |
| **Oxford** | SCID interview | MDD subjects: psychosis or substance dependence (DSM-IV), clinically significant risk of suicidal behaviour, having contraindications to escitalopram treatment or being treated with psychotropic medication less than three weeks before the study (five weeks in the case of fluoxetine);<br>Control subjects: current or past history of Axis I disorder as defined by DSM-IV; Both groups: major somatic or neurological disorders, pregnancy or breast-feeding, contraindications to MR imaging or concurrent medication which could alter emotional processing |
| **FOR2107** | SCID-1 | **Marbourg:** |

| | | |
|---|---|---|
| | | All subjects: Exclusion criteria all: any MRI contraindications; any neurological abnormalities.<br>Control subjects: any current or former psychiatric disorder;<br>MDD subjects: substance dependence or current benzodiazepine treatment (wash out of at least three half-lives before study participation)" |
| | | **Münster**<br>All subjects: any MRI contraindications; any neurological abnormalities. Exclusion criteria controls: any current or former psychiatric disorder; Exclusion criteria patients: substance dependence or current benzodiazepine treatment (wash out of at least three half-lives before study participation)" |
| **AFFDIS** | ICD-10/Dsm IV criteria | All subjects: current or history of neurological disorder or brain injury, current substance abuse or dependence (not including nicotine), pregnancy, MRI contraindications, inability to give consent.<br>MDD subjects: comorbid psychiatric diagnosis.<br>Control subjects: current or history of psychiatric diagnosis. |
| **Singapore** | SCID interview | All subjects: Exclusion criteria 1) History of significant head injury 2) Neurological diseases such as epilepsy, cerebrovascular accident 3) Impaired thyroid function 4) Steroid use 5) DSM IV alcohol or substance use or dependence 6) Contraindications to MRI (e.g. pacemaker, orbital foreign body, recent surgery/procedure with metallic devices/implants deployed) using standard MRI Request Form from NNI 7) Pregnant women 8) Claustrophobia |
| **ETPB** | HAMD,BDI, SHAPS,MADRS | Current or past diagnosis of Schizophrenia or any other psychotic disorder as defined in the DSM-IV. Subjects with a history of DSM-IV drug or alcohol dependency or abuse (except for nicotine or caffeine) within the preceding 3 months. Female subjects who are either pregnant or nursing. Serious, unstable illnesses including hepatic, renal, gastroenterological, respiratory, cardiovascular (including ischemic heart disease), endocrinological, neurologic, immunologic, or hematologic disease. Subjects with uncorrected hypothyroidism or hyperthyroidism. Subjects with one or more seizures without a clear and resolved aetiology. Treatment with a reversible MAOI within 4 weeks prior to study phase I. Treatment with fluoxetine within 5 weeks prior to study phase I. Treatment with any other concomitant medication not allowed (Appendix A for Sub study 2; Appendix G for Sub study 4) 14 days prior to study phase I. No structured psychotherapy will be permitted during the study. Current NIMH employee/staff or their immediate family member.<br>MDD subjects: Previous treatment with ketamine or hypersensitivity to amantadine. Additional Exclusion Criteria for Sub study 4 (patients with MDD or BD). Subjects who currently are using drugs (except for caffeine or nicotine), must not have used illicit substances in the 2 weeks prior to screen and must have a negative alcohol and drug urine test (except for prescribed benzodiazepines) urine test at screening. Presence of any medical illness likely to alter brain morphology and/or physiology (e.g., hypertension, diabetes) even if controlled by medications. Clinically significant abnormal laboratory tests. Presence of metallic (ferromagnetic) implants (e.g, heart pacemaker, aneurysm clip). Subjects who, in the investigator s judgment, pose a current serious suicidal or homicidal risk, or who have a MADRS item 10 score of >4. |
| **BiDirect** | M.I.N.I. Neuropsychiatric Interview, IDS, HAMD, CESD, ICD-10 | All subjects: dementia, addiction |
| **Sydney** | SCID interview | MDD subjects: presence of axis-I disorders other than MDD, panic disorder, social anxiety disorder, or generalized anxiety disorder. Control subjects: no Axis-I diagnosis, no medication use. Exclusion criteria for all subjects included medical instability (as determined by a psychiatrist), history of neurological disease (e.g. tumour, head trauma, epilepsy), medical illness known to impact cognitive and brain function (e.g. cancer), intellectual and/or developmental disability and insufficient English for neuropsychological assessment. All subjects were asked to abstain from drug or alcohol use for 48 hours prior to testing and informed about a drug screen protocol. |
| **Moral Dilemma** | SCID interview | MDD subjects: lifetime or current SCID-I diagnosis of psychotic disorder, or bipolar I or II disorder; current antidepressant medication use.<br>Control subjects: any SCID-I diagnosis or medication use. Both groups: Acute or unstable medical disorder; general MRI contraindications |

| | | |
|---|---|---|
| **Stanford FAA** | SCID interview | MDD subjects: presence of axis-I disorders other than MDD, anxiety and eating disorders<br>Control subjects: control individuals did not meet diagnostic criteria for any current psychiatric. Both groups: alcohol / substance abuse or dependence within six months prior to MRI scanning, history of head trauma with loss of consciousness > 5 min, aneurysm, or any neurological or metabolic disorders that require ongoing medication or that may affect the central nervous system (including thyroid disease, diabetes, epilepsy or other seizures, or multiple sclerosis), MRI contraindications, or bad MRI data (e.g., extreme movement). |
| **FIDMAG** | DSM-IV-TR criteria | MDD subjects: excluded (i) if they were left-handed; (ii) if they were younger than 18 or older than 65 years; (iii) if they had a history of brain trauma or neurological disease; (iv) if they had shown alcohol/ substance abuse within 12 months prior to participation; and (v) if they had undergone electroconvulsive therapy in the previous 12 months. |
| **SoCAT** | SCID interview | MDD subjects: 1) History of significant head injury 2) Neurological diseases such as epilepsy, cerebrovascular accident 3) Other diagnoses on Axis I disorders4) |
| **SHIP-TREND-0** | M-CIDI interview | MDD subjects: no special exclusion criteria Control subjects: no lifetime diagnosis of depression, no antidepressants, and severity index=0<br>All subjects: We removed subjects with due to medical conditions (e.g. a history of cerebral tumour, stroke, Parkinson's diseases, multiple sclerosis, epilepsy, hydrocephalus, enlarged ventricles, pathological lesions) or due to technical reasons (e.g. severe movement artefacts or inhomogeneity of the magnetic field). |
| **Hiroshima** | MINI | MDD patients: comorbid psychiatric disorders other than MDD<br>Control subjects: any history of psychiatric disorder |
| **CSAN** | MINI | MDD subjects: a current DSM-5 diagnosis of substance use disorder, except nicotine; a psychotic disorder, except depression with mood-congruent psychotic features; new antidepressant medication during the month before study participation (two months for fluoxetine); change of the dose of psychotropic medications over the last month (antidepressant and antipsychotic medication) or the last two months (mood stabilizers and anticonvulsants). |
| **SHIP-2** | M-CIDI interview | MDD subjects: presence of axis-I disorders other than MDD, anxiety disorders, conversion, somatization and eating disorder.<br>Control subjects: no lifetime diagnosis of depression, no anti-depressiva, and severity index=0 All subjects: We removed subjects with medical conditions (e.g. a history of cerebral tumour, stroke, Parkinson's diseases, multiple sclerosis, epilepsy, hydrocephalus, enlarged ventricles, pathological lesions) or due to technical reasons (e.g. severe movement artefacts or inhomogeneity of the magnetic field). |
| **MPIP** | M-CIDI/SCAN interview | MDD subjects: Munich Antidepressant Response Signature (MARS) study MDD subjects (clinical consensus diagnosis or M-CIDI (since 2008)): depressive syndromes secondary to any medical or neurological condition (e. g., intoxication, drug abuse, stroke), the presence of manic, hypomanic or mixed affective symptoms, lifetime diagnosis of alcohol dependence, illicit drug abuse or the presence of severe medical conditions (e.g., ischemic heart disease). Patients with bipolar depression were excluded for the current MR study. Control subjects: age > 65, MMSE<27, presence of severe somatic diseases or lifetime history of the following axis I disorders as assessed by the M-CIDI interview: alcohol dependence, drug abuse or dependence, possible psychotic disorder, mood disorder, anxiety disorder including OCD and PTSD, somatoform disorder, dissociative disorder NOS, and eating disorder 2. Recurrent unipolar depression (RUD) study: MDD subjects (SCAN interview): presence of manic episodes, mood incongruent psychotic symptoms, the presence of a lifetime diagnosis of intravenous drug abuse and depressive symptoms only secondary to alcohol or substance abuse or to medical illness or medication.<br>Control subjects: presence of severe somatic diseases or life-time history of anxiety and affective disorders according to the Composite International Diagnostic-Screener (CIDI-S). All subjects: gross incidental MR findings such as territorial infarction, tumour, hydrocephalus, malformations and anatomical deviations (e.g. enlarged ventricles) that prevent appropriate image processing were additional exclusion criteria. 3. MR images of 9 additional controls acquired at the LMU, Munich, meeting equivalent criteria as the RUD control sample were included. |

MDD: Major Depressive Disorder; CIDI: the Composite International Diagnostic Interview; SCID: Structured Clinical Interview for DSM disorders; SCAN: Schedules for Clinical Assessment in Neuropsychiatry; MINI: M.I.N.I. International Neuropsychiatric Interview; CESD: Center for Epidemiologic Studies Depression scale; DSM: Diagnostic and Statistical Manual of Mental Disorders; MRI: Magnetic Resonance Imaging; OCD: Obsessive Compulsive Disorder; PTSD: Posttraumatic Stress Disorder; HAMD: Hamilton Depression Scale; BDI: Beck Depression Inventory; SHAPS: Snaith-Hamilton pleasure scale; MARDS: Montgomery-Asberg Depression Rating Scale; KSDAS; CDRS-R: Children's Depression Rating Scale–Revised; SADS: Schedule for Affective Disorders and Schizophrenia for School-Age Children–Present and Lifetime

#### S1.3 ENIGMA - Major Depressive Disorder Working Group image acquisition criteria, breakdown by site

| Cohort | Scanner type | Sequence T1 | FS version | Slice Orient. | OS |
|---|---|---|---|---|---|
| **ClinG** | 3T Siemens Tim Trio | Standard 3D T1-weighted turbo fast low angle shot (turbo FLASH); voxel size 1 mm x 1 mm x 1mm (based on the ADNI protocol (Jack et al. 2008); TR=225 msec; TE=3.26 msec, FOV=256 x 256 x 192 | 5.3 | Sagittal | Linux |
| **Edinburgh (Bipolar Family Study)** | 1.5T GE Signa | T1-weighted sequence. TR=500 msec; TE=4 msec; flip angle 8°; matrix 192 x 192; 180 slices; voxel size 1.25 mm x 1.25 mm x 1.20 mm; FOV=24, phase FOV 1 | 5.3 | Coronal | linux 6, x86_64, kernel 2.6.32 |
| **McMaster University Mood Disorders** | 1.5T (GE); 3T(GE) | 1.5-T. Sigma GE Genesis-based Echo-Speed scanner running version 5.7 software and using a standard 30-cm circularly polarized head coil. Sagittal anatomic images were acquired by using a 3D/FSPGR/20 sequence (flip angle=20; echo delay time in-phase (TE), minimum repetition time (TR)=300 ms; inversion recovery=300 ms; matrix=512x256; field of view (FOV)=24 cm; scan thickness=1.2 mm). 3-T MRI Sigma GE Genesis (General Electric Medical Systems, Milwaukee, WI). Sagittal T-1 weighted images were acquired using a 3D FSPGR-IR sequence, (TR/TE=10.3/2.1 ms; flip angle=20; inversion time=300; matrix=512x256; FOV=24; and slice thickness=1.2 mm. | 5 | | |
| **Houston** | 1.5 T Philips Medical Systems Gyroscan Intera; | Subjects in the 20000s: Fast field echo sequence- repetition time (TR) = 24 ms, echo time (TE) = 4.99 ms, flip angle = 40°, slice thickness = 1 mm, matrix size = 256 × 256 and 150 slices. Subjects in 30000s: MPRAGE- repetition time (TR) = 1750 ms, echo time (TE) = 4.39 ms, flip angle = 8°, slice thickness = 1 mm, matrix size = 208 × 256 and 160 slices. | 5.3 | Sagittal; | Fedora 19 |
| | 3T Siemens Allegra | | | Transverse | |
| **Münster Neuroimaging Cohort** | 3T Philips Gyroscan Intera | 3D fast gradient echo sequence (turbo field echo), repetition time = 7.4 milliseconds, echo time = 3.4 milliseconds, flip angle = 9°, two signal averages, inversion prepulse every 814.5 milliseconds, acquired over a field of view of 256 (feet - head [FH]) × 204 (anterior -posterior [AP]) × 160 (right -left [RL]) mm, phase encoding in AP and RL direction, reconstructed to cubic voxels of .5 mm × .5 mm × .5 mm | 5.3 | Sagittal | Red Hat Enterprise Linux Server release 5.11 (Tikanga) |
| **Melbourne** | 3T GE Signa Excite | 3D BRAVO sequence 140; TR=7900 ms; TE=3000 ms; flip angle=13º; FOV=256 mm; matrix=256 x 256 | 5.3 | Axial | Linux Debian x86 64 |
| **Stanford T1w Aggregate** | 1.5T GE Signa Excite | Whole-brain T1-weighted images were collected using a spoiled gradient echo (SPGR) pulse sequence (116 sagittal slices; through-plane resolution = 1.5 mm; in-plane resolution = 0.86 x 0.86 mm; flip angle = 15 degrees; repetition time | 5.3 | Sagittal | Linux-centos6_x86_64 |

| | | | | | |
|---|---|---|---|---|---|
| | | [TR] = 8.3-10.1 ms; echo time [TE] = 1.7-3.0; inversion time [TI] = 300 ms; matrix = 256 x 192). | | | |
| **Magdeburg-** sexpect | 3 Tesla Siemens MAGNETOM Trio scanner (Siemens, Erlangen, Germany) | High resolution T1 -weighted structural MRI scans of the brain were acquired for structural reference using a 3D - MPRAGE sequence (TE = 4.77 ms, TR = 2500 ms, T1 = 1100 ms, flip angle = 7°, bandwidth = 140 Hz/pixel, acquisition matrix = 256 × 256 × 192, isometric voxel size = 1.0 mm3). | 5.3 | Sagittal | Oracle Linux Server_x86_64 |
| **BRCDECC London** | 1.5T GE Signa HDx | ADNI-1 MPRAGE pulse sequence (details at http://adni.loni.ucla.edu/research/protocols/mri-protocols/) | 5.3 | Sagittal | Linux-centos4_x86_64 |
| **Barcelona** | 3T Philips Achieva | 3D MPRAGE images (Whole-brain T1-weighted); TR=6.7ms, TE=3.2ms; 170 slices, voxel size 0.89X0.89X1.2 mm. Image dimensions 288X288X170; field of view: 256X256X204; slice thickness: 1.2 mm; with a sagittal slice orientation, T1 contrast enhancement, flip angle: 8º, grey matter as a reference tissue, ACQ matrix MXP = 256X240 and turbo-field echo shots (TFE) = 218. | 6 | Sagittal | Scientific Linux 5 |
| **Houston adolescents** | .5 T Philips Medical Systems Gyroscan Intera;<br>3T Siemens Allegra | Subjects in the 20000s: Fast field echo sequence- repetition time (TR) = 24 ms, echo time (TE) = 4.99 ms, flip angle = 40°, slice thickness = 1 mm, matrix size = 256 × 256 and 150 slices. Subjects in 30000s: MPRAGE- repetition time (TR) = 1750 ms, echo time (TE) = 4.39 ms, flip angle = 8°, slice thickness = 1 mm, matrix size = 208 × 256 and 160 slices. | 5.3 | Sagittal;<br>Transverse | Fedora 19 |
| **EPISCA (Leiden)** | 3T Philips Achieva | a sagittal 3-dimensional gradient-echo T1-weighted image was acquired (repetition time = 9.8 ms; echo time = 4.6 ms; flip angle = 8°; 140 sagittal slices; no slice gap; field of view =256 × 256 mm; 1.17 × 1.17 × 1.2 mm voxels; duration = 4:56 min) | 5.3 | Sagittal | Ubuntu 14.04.5 LTS (Linux 3.13.0-153-generic x86_64) |
| **UCSF** | 3T GE Discovery MR750 | SPGR T1-weighted: TR=8.1 ms; TE=3.17 ms; TI=450 ms; flip angle=12°; 256x256 matrix; FOV=250x250 mm; 168 sagittal slices; slice thickness=1 mm; in-plane resolution=0.98x 0.98 mm | 5.3 | Sagittal | Linux-centos6_x86_64-stable-pub-v5.3.0. |
| **Sao Paulo (Wellcome)** | 1.5T General Eletric (GE) | Imaging data were acquired using two MRI scanners (at the Clinics Hospital of the University of Sao Paulo 1.5 T GE Signa scanner, General Electric, Milwaukee Wisconsin, USA). T1-SPGR sequence providing 124 contiguous slices, voxel size 0.8660.8661.5 mm, echo time 5.2 ms, resolution time 21.7 ms, flip angle 20, field of vision 22, matrix 256x192) | 5.3 | | |
| **Minnesota** | 3.0 Tesla Tim Trio scanner; Siemens Corp | A 5-minute structural scan was acquired using a T1-weighted, high-resolution, magnetization-prepared gradient-echo sequence: repetition time, 2530 milliseconds; echo time, 3.65 milliseconds; inversion time, 1100 milliseconds; | 5.3 | Coronal | Linux |

| | | | | | |
|---|---|---|---|---|---|
| | | flip angle, 7°; field of view, 256 × 176 mm; voxel size, 1-mm isotropic; 224 slices; and generalized, autocalibrating, partially parallel acquisition acceleration factor, 2. | | | |
| **Calgary** | 1.5T Siemens Magnetom Vision. 3T GE Discovery MR750 | 1.5T: A sagittal scout series was acquired to test image quality. 3D fast low angle shot (FLASH) sequence was used to acquire data from 124 1.5 mm-thick contiguous coronal slices through the entire brain (echo time = 5ms, repetition time = 25ms, acquisition matrix = 256 x 256 pixels, field of view = 24 cm and flip angle = 40°). 3T: Anatomical imaging acquisition parameters: axial acquisition, repetition time (TR), 2200 milliseconds (ms); echo time (TE), 3.04 ms; TI, 766, 780; flip angle, 13 degrees; 208 partitions; 256 × 256 matrix; and field of view, 256. | 5.3 | coronal;, axial | MacOs Sierra |
| **QTIM** | Bruker 4T Wholebody MRI | 3D T1 weighted sequence. TR=1500 msec; TE=3.35 msec; flip angle=8°, 256 or 240 (coronal or sagittal) slices, FOV=240 mm, matrix 256x256x256 (or 256x256x240) | 5.1 | Coronal, then sagittal following software upgrade. | Linux- centos4_x86_64-stable-pub-v5.1.0 |
| **Oxford** | 3T Siemens Tim Trio | Voxel resolution 0.78 x 0.8 x 0.78 mm on a 208 x 256 x 200 grid, TE/TI/TR= 4.8/1100/2040 ms | 5.3 | | |
| **FOR2107** | **Marbourg:** 3T Siemens Magnetom TiroTim syngo MR B17<br><br>**Münster:** 3T Siemens PRISMA | **Marbourg:** Sequence: 3D T1-weighted magnetization prepared rapid acquisition gradient echo (MPRAGE) - Sagittal Acquisition Direction, # of Slices 176, 0.5mm Slice Gap, 1.0x1.0x1.0 Voxel Size (mm3), TI 900 ms, TE 2.26 ms, TR 1900 ms, Flip Angle 9.<br>**Münster:** Sequence: 3D T1-weighted magnetization prepared rapid acquisition gradient echo (MPRAGE). - Sagittal Acquisition Direction, # of Slices 192, 0mm Slice Gap, 1.0x1.0x1.0 Voxel Size (mm3), TI 900 ms, TE 2.28 ms, TR 1900 ms, Flip Angle 8 | 5.3 | Sagittal | Red Hat Enterprise Linux Server release 5.11 (Tikanga) |
| **AFFDIS** | 3T Siemens Magnetom TrioTim | 3D T1 (176 slices; TR = 2250 ms; TE = 3.26 ms; FOV 256; voxel size 1X1X1mm) | 5.3 | Sagittal | Linux CentOS |
| **Singapore** | Achieva 3T, Philips Medical Systems, Netherlands | Whole brain high resolution 3D MP-RAGE (magnetisation-prepared rapid acquisition with a gradient echo) volumetric scans (TR/TE/TI/flip angle 8.4/3.8/3000/8; matrix 256x204; FOV 240mm2) with axial orientation (reformatted to coronal) | 5.3 | Axial | Linux_Ubuntu12.04_6 4 |
| **ETPB** | 3T, GE HDx | Fast spoiled gradient recalled echo (FSPGR). Slice Thickness: 1. Repetition Time: 8.836. Echo Time: 3.496. Inversion Time: 450. Magnetic Field Strength: 3. Spacing Between Slices: 1. Echo Train Length: 1. Percent Sampling: 100. Percent Phase Field of View: 100. Pixel Bandwidth: 195.312. Reconstruction Diameter: 256. Acquisition Matrix: | 5.3 | Sagittal | Linux |

| | | | | | |
|---|---|---|---|---|---|
| | | 0,256,256,0. In-plane Phase Encoding Direction: ROW. Flip Angle: 13 | | | |
| **BiDirect** | 3 T Philips Intera scanner | 3D T1-weighted turbo field echo images were collected with a the following parameters: TR = 7.26, TE = 3.56, 9° flip angle, 160 sagittal slices, matrix dimension 256 x 256, FOV = 256 x 256mm, 2mm slice thickness (reconstructed to 1mm) and a resulting voxel size of 1x1x1mm | 6 | Sagittal | freesurfer-Linux-centos6_x86_64-dev-20161222-499fc91 |
| **Sydney** | 3T GE MR750 | 3D T1-weighted sequence. TR=7.2 msec; TE=2.78 msec; matrix =256; FOV=240; No. slices=196; thick=0.9mm; inplane resolution=0.9375 | 5.1 | Coronal | Linux_Ubuntu16.04 lts 64bit |
| **Moral Dilemma** | 3T GE Signa Excite | 3D BRAVO sequence: 140 contiguous slices; repetition time, 7900 ms; echo time, 3000 ms; flip angle, 13°; in a 25.6-cm field of view, with a 256 × 256 pixel matrix and a slice thickness of 1 mm (1 mm gap). | 5.3 | Axial | Linux Debian x86 64 |
| **Stanford FAA** | 3.0T GE Discovery MR750 | Whole-brain T1-weighted images were collected using a spoiled gradient echo (SPGR) pulse sequence (186 sagittal slices; resolution = 0.9 mm isotropic; flip angle = 12°; repetition time [TR] = 6,240 ms; echo time [TE] = 2.34 ms) | 5.3 | Sagittal | Centos6_x86_64, Linux-based HPC |
| **FIDMAG** | 1.5T, GE Signa | 3D T1: matrix size = 512 × 512, 180 contiguous axial slices, voxel resolution = 0.47 × 0.47 × 1mm, no slice gap, TE = 3.93ms, TR = 2000ms and inversion time (TI) = 710ms, flip angle = 15 degrees | 6 | Axial | Linux-centos6_x86_64 |
| **SoCAT** | 3.0 T, Siemens Verio,Numaris/4,Syngo MR B17,Erlangen,Germany | 3D T1 weighted MP-Rage/axial plane; TR=1900 msec; TE=3.4 msec; Flip angle=15°; Voxel size 1 mm x 1 mm x 1 mm | 7 | Axial | Ubuntu 18.04 LTS |
| **SHIP-TREND-0** | 1.5T Siemens Avanto | 3D T1-weighted (MP-RAGE/ axial plane); TR=1900 msec; TE=3.4 msec; Flip angle=15°; voxel size 1 mm x 1 mm x 1 mm | 5.3 (cortical), 5.1 (subcortical) | Axial | Centos6_x86_64 |
| **Hiroshima** | 3T Siemens (Spectra, Verio.Dot), 3T GE (Signa HDxt) Site 1 = GE Signa HDxt 3.0T<br>2= GE Signa HDxt 3.0T<br>3 = SIEMENS MAGNETOM Spectra 3.0T<br>4 = SIEMENS MAGNETOM Verio.Dot 3.0T | T1 256x256x256 matrix of 1x1x1mm voxels (Siemens: ADNI MPRAGE (tfl), GRAPPA, 192 slices, GE: SPGR, 184 slices) *Detailed scanning parameter sheets are available for all 4 scanners on request. | 5.3 | Sagittal | Linux_Ubuntu_18.04 |
| **CSAN** | 3T Siemens MAGNETOM PRISMA | Whole-head t1-weighted MPRAGE (TR = 2300 ms, TE = 2.34 ms, FOV 250 × 250 mm, voxel size = 0.9 × 0.868 × 0.868 mm, flip angle = 8°). | 7.2 | Sagittal | Ubuntu |

| | | | | | |
|---|---|---|---|---|---|
| **SHIP** | 1.5T Siemens Avanto | 3D T1-weighted (MP-RAGE/ axial plane); TR=1900 msec; TE=3.4 msec; Flip angle=15°; voxel size 1 mm x 1 mm x 1 mm | 5.3 (cortical), 5.1 (subcortical) | Axial | Centos6_x86_64 |
| **MPIP** | 1.5T GE and Siemens | #1: T1-weighted SPGR sagittal 3D volume. TR=1030 msec; TE=3.4 msec; 124 slices; matrix=256x256; FOV=23.0x23.0 cm2; voxel size=0.8975 mm x0.8975 mm x 1.2- 1.4 mm; flip angle=90°; birdcage resonator. #2: same scanner as #1, platform update Signa Excite, sagittal T1-weighted (spin echo sequence, TR=9.7 msec, TE=2.1 msec; FOV=25.0x25.0 cm2, voxel size=0.875 mm x0.875 mm x1.2 mm, 124- 132 slices, flip angle=90°. #3: Siemens 1.5 Tesla, Vario, 3D MPRAGE, TR=11.6 msec; TE=4.9 msec; FOV 23x23 cm2; matrix 512x512; 126 axial slices; voxel site 0.45 mm x 0.45 mm x 1.5 mm. (only N=2 subjects) | 5.3 | 1.5 GE: sagittal. 1.5 Siemens: axial | Linux 2.6.37.1-1.2-desktop x86_64 |

**S1.4: Age of onset, BDI and HDRS values of individuals with depression per site. Not all sites had respective information available.**

| | MDD | | | | | |
|---|---|---|---|---|---|---|
| Site name | mean AO (SD) | | mean BDI (SD) | | mean HDRS (SD) | |
| ClinG | 30.4 | 10.6 | 21.9 | 9.8 | 20.0 | 4.3 |
| Edinburgh (Bipolar Family Study) | NA | NA | NA | NA | 5.1 | 5.4 |
| McMaster University Mood Disorders Houston | 22.5 | 11.0 | NA | NA | 11.9 | 7.6 |
| Houston | 22.7 | 14.0 | 16.5 | 15.1 | 9.9 | 7.9 |
| Münster Neuroimaging Cohort | 29.5 | 11.9 | 25.9 | 10.1 | 19.3 | 4.2 |
| Melbourne | 15.9 | 2.7 | NA | NA | NA | NA |
| Stanford T1w Aggregate | 19.5 | 9.8 | 26.5 | 10.3 | 15.1 | 5.7 |
| Magdeburg - Sexpect | 26.2 | 7.5 | 21.0 | 9.5 | 12.4 | 3.0 |
| BRCDECC London | 20.4 | 9.3 | 15.3 | 11.4 | NA | NA |
| Barcelona | 33.2 | 11.4 | NA | NA | 13.7 | 8.2 |
| Houston adolescents | 6.3 | 5.7 | NA | NA | 11.0 | 6.4 |
| UCSF | 13.3 | 2.3 | 26.7 | 11.9 | NA | NA |
| Sao Paulo (Wellcome) | NA | NA | NA | NA | 15.4 | 9.5 |
| Minnesota | 12.4 | 2.4 | 25.8 | 12.1 | NA | NA |
| Calgary | 14.3 | 3.2 | 26.8 | 11.6 | 19.1 | 6.6 |
| QTIM | 18.4 | 3.3 | NA | NA | NA | NA |
| Oxford | 25.6 | 9.1 | NA | NA | 22.9 | 4.3 |
| FOR2107 | 26.6 | 12.7 | 18.2 | 11.1 | 8.7 | 6.8 |
| AFFDIS | 31.0 | 15.6 | 28.9 | 11.2 | 23.4 | 9.8 |
| Singapore | 34.4 | 8.7 | NA | NA | 5.6 | 5.3 |
| ETPB | 15.8 | 6.6 | 29.3 | 7.5 | 21.6 | 4.3 |
| BiDirect | 38.5 | 10.8 | NA | NA | 13.6 | 6.7 |
| Sydney | 24.2 | 17.0 | NA | NA | 12.3 | 6.9 |
| StanfFAA | 16.3 | 6.8 | 28.3 | 10.0 | 18.9 | 4.2 |
| FIDMAG | 37.2 | 13.4 | NA | NA | 24.8 | 5.8 |
| Socat | 31.6 | 16.7 | 24.0 | 12.1 | 13.3 | 7.6 |
| SHIP-TREND-0 | 36.2 | 14.3 | 12.4 | 8.1 | NA | NA |
| Hiroshima | 38.2 | 13.3 | 29.8 | 9.4 | 18.7 | 5.6 |
| SHIP- | 38.4 | 13.2 | 11.7 | 10.4 | NA | NA |
| MPIP | 35.1 | 14.1 | 14.1 | 11.0 | 24.9 | 6.6 |

**AO: Age of onset. BDI: Beck depression inventory. HDRS: Hamilton Depression Rating Scale**

### S1.5: Percentage of MDD patients using antidepressant medication, percentage of first episode and recurrent episode MDD patients, percentage of acutely depressed and remitted MDD patients, break down per site.

| Site | Acutely depressed | remit ted | first episode | recurrent episode | antidepressa nt free | anti-depressant use |
|---|---|---|---|---|---|---|
| **CLING** | 3 | 46 | 23 | 26 | 3 | 46 |
| **Edinburgh (Bipolar Family Study)** | 0 | 18 | 0 | 0 | 3 | 0 |
| **McMaster University Mood Disorders Houston** | 0 | 51 | 22 | 29 | 22 | 29 |
| **Houston** | 39 | 37 | 21 | 43 | 74 | 0 |
| **Münster Neuroimaging Cohort** | 16 | 215 | 49 | 181 | 22 | 193 |
| **Melbourne** | 0 | 142 | 48 | 88 | 120 | 22 |
| **Stanford T1w Aggregate** | 0 | 48 | 6 | 40 | 22 | 19 |
| **Magdeburg - Sexpect** | 0 | 5 | 1 | 4 | 0 | 5 |
| **BRCDECC London** | NA | NA | 0 | 69 | 19 | 50 |
| **Barcelona** | 23 | 39 | 22 | 40 | 4 | 58 |
| **Houston adolescents** | NA | NA | 20 | 7 | 27 | 1 |
| **Episca (Leiden)** | 0 | 19 | 19 | 0 | 18 | 1 |
| **UCSF** | 6 | 59 | 31 | 34 | 74 | 0 |
| **Sao Paulo (Wellcome)** | 0 | 18 | 5 | 11 | 11 | 13 |
| **Minnesota** | 6 | 0 | 16 | 22 | 52 | 16 |
| **Calgary** | 0 | 37 | 0 | 37 | 20 | 17 |
| **Calgary** | 0 | 18 | 18 | 0 | 17 | 1 |
| **QTIM** | NA | NA | NA | NA | 73 | 29 |
| **Oxford** | 0 | 38 | 19 | 19 | 38 | 0 |
| **FOR2107** | 73 | 234 | 91 | 183 | 117 | 190 |
| **FOR2107** | 52 | 111 | 58 | 102 | 65 | 98 |
| **AFFDIS** | 0 | 28 | 2 | 26 | 1 | 27 |
| **Singapore** | NA | NA | 8 | 14 | 4 | 18 |
| **ETPB** | 0 | 34 | 0 | 34 | 34 | 0 |
| **BiDirect** | 0 | 572 | 323 | 249 | 71 | 501 |
| **Sydney** | 164 | 37 | 58 | 151 | 83 | 128 |
| **Moral Dilemma** | 0 | 24 | 8 | 16 | 24 | 0 |
| **StanfFAA** | 0 | 14 | 0 | 14 | 11 | 3 |
| **FIDMAG** | 1 | 33 | 10 | 22 | 3 | 30 |
| **Socat** | 33 | 46 | 19 | 60 | 41 | 38 |
| **SHIPtrend-0** | NA | NA | 113 | 199 | 258 | 54 |
| **Hiroshima** | NA | 150 | 74 | 74 | 10 | 138 |
| **CSAN** | NA | NA | NA | NA | NA | NA |
| **SHIP** | NA | NA | 75 | 60 | 111 | 24 |
| **MPIP** | 46 | 291 | 91 | 246 | 53 | 284 |

**NA: not provided**

#### S1.6 ENIGMA - Major Depressive Disorder Working Group Clinical characteristics that were shared across individuals with MDD and healthy controls. BMI (Body Mass Index), CTQ (Children Trauma Score) for individuals with depression and healthy controls breakdown for available participating sites.

| | HC (Train and test set) | | | | MDD | | | |
|---|---|---|---|---|---|---|---|---|
| **Site name** | mean BMI | (SD) | mean CTQ | (SD) | mean BMI | (SD) | mean CTQ | (SD) |
| **Münster Neuroimaging Cohort** | 24.6 | 4.1 | 33.5 | 8.3 | 25.6 | 3.2 | 45.6 | 16.6 |
| **Melbourne** | NA | NA | NA | NA | 25.9 | 6.5 | NA | NA |
| **LOND** | 24.0 | 2.9 | 34.8 | 13.0 | 26.1 | 5.1 | 41.9 | 12.5 |
| **Houston adolescents** | 22.8 | 6.1 | NA | NA | 24.7 | 5.7 | NA | NA |
| **UCSF** | NA | NA | 31.8 | 7.6 | NA | NA | 57.4 | 16.1 |
| **Minnesota** | 24.1 | 5.6 | NA | NA | 24.3 | 6.2 | NA | NA |
| **Calgary** | 20.8 | 4.7 | NA | NA | 24.4 | 4.9 | NA | NA |
| **QTIM** | 23.1 | 3.8 | NA | NA | 23.7 | 5.1 | NA | NA |
| **FOR2107** | NA | NA | 32.2 | 8.2 | NA | NA | 45.9 | 15.8 |
| **AFFDIS** | NA | NA | 34.8 | 11.2 | NA | NA | 46.1 | 14.4 |
| **Singapore** | 26.6 | 5.5 | NA | NA | 24.4 | 4.9 | NA | NA |
| **ETPB** | 27.8 | 4.7 | NA | NA | 27.6 | 6.8 | NA | NA |
| **Sydney** | 22.8 | 3.1 | NA | NA | 23.2 | 4.2 | 45.4 | 14.8 |
| **Stanford FAA** | NA | NA | 33.4 | 8.4 | NA | NA | 49.4 | 23.3 |
| **SoCAT** | NA | NA | 27.9 | 3.2 | NA | NA | 46.0 | 15.5 |
| **SHIP-TREND-0** | 27.4 | 4.2 | 31.4 | 7.1 | 27.7 | 4.7 | 36.8 | 13.7 |
| **SHIP** | 27.7 | 4.2 | 32.2 | 7.4 | 27.4 | 5.3 | 37.1 | 10.7 |
| **MPIP** | NA | NA | NA | NA | NA | NA | 77.0 | NA |

#### S1.7: Z-score group differences between HC and individuals with MDD. Cohen's d; p-values of the difference (FDR corrected); distributional overlap score (range: 0: no overlap; 1: complete overlap), breakdown per region.

| Region of Interest | Cohen's d | Adjusted p-value | Distributional overlap score |
|---|---|---|---|
| Banks of the Superior Temporal Sulcus | 0.117 | **0.0001** | 0.935 |
| Caudal anterior cingulate cortex | 0.053 | 0.0728 | 0.975 |
| Caudal middle frontal gyrus | 0.036 | 0.2067 | 0.964 |
| Cuneus | 0.005 | 0.8599 | 0.977 |
| Entorhinal cortex | 0.049 | 0.0915 | 0.958 |
| Frontal pole | 0.026 | 0.3621 | 0.975 |
| Fusiform gyrus | 0.169 | **0.0000** | 0.923 |
| Inferior parietal gyrus | 0.109 | **0.0002** | 0.950 |
| Inferior temporal gyrus | 0.126 | **0.0001** | 0.947 |
| Insula | 0.116 | **0.0001** | 0.939 |
| Isthmus cingulate cortex | 0.059 | **0.0474** | 0.964 |
| Lateral occipital gyrus | 0.060 | **0.0458** | 0.972 |
| Lateral orbitofrontal cortex | 0.071 | **0.0223** | 0.957 |
| Lingual gyrus | 0.046 | 0.1165 | 0.959 |
| Medial orbitofrontal cortex | 0.117 | **0.0001** | 0.935 |
| Middle temporal gyrus | 0.126 | **0.0001** | 0.944 |
| Paracentral gyrus | 0.097 | **0.0011** | 0.961 |
| Parahippocampal gyrus | 0.069 | **0.0237** | 0.969 |
| Pars opercularis of the inferior frontal gyrus | 0.106 | **0.0003** | 0.933 |
| Pars orbitalis of the inferior frontal gyrus | 0.070 | **0.0223** | 0.965 |
| Pars triangularis of the inferior frontal gyrus | 0.058 | **0.0480** | 0.971 |
| Pericalcarine gyrus | -0.040 | 0.1704 | 0.953 |
| Postcentral gyrus | 0.065 | **0.0290** | 0.967 |
| Posterior cingulate cortex | 0.076 | **0.0129** | 0.967 |
| Precentral gyrus | 0.110 | **0.0002** | 0.935 |
| Precuneus | 0.068 | **0.0244** | 0.951 |
| Rostral anterior cingulate cortex | 0.110 | **0.0002** | 0.943 |
| Rostral middle frontal gyrus | 0.055 | 0.0622 | 0.958 |
| Superior frontal gyrus | 0.060 | **0.0458** | 0.959 |
| Superior parietal gyrus | 0.043 | 0.1406 | 0.965 |
| Superior temporal gyrus | 0.096 | **0.0012** | 0.959 |
| Supramarginal gyrus | 0.109 | **0.0002** | 0.945 |
| Temporal pole | 0.019 | 0.5127 | 0.984 |
| Transversetemporal gyrus | 0.067 | **0.0271** | 0.957 |
| Average cortical thickness | 0.112 | **0.0002** | 0.934 |

Cohen's d; p-values of the difference (FDR corrected); distributional overlap score (range: 0: no overlap; 1: complete overlap), breakdown per region. Significant differences ($p < 0.005$) are marked in bold

#### S1.8 Percentage overlap: Percentage overlap per region.

| % Overlap negative deviations, HC | % Overlap positive deviations, HC | % Overlap negative deviations, MDD | % Overlap positive deviations, MDD | Region |
|---|---|---|---|---|
| 0.089 | 0.092 | 0.094 | 0.072 | Banks of the Superior Temporal Sulcus |
| 0.081 | 0.094 | 0.069 | 0.091 | Caudal anterior cingulate cortex |
| 0.100 | 0.079 | 0.070 | 0.077 | Caudal middle frontal gyrus |
| 0.054 | 0.116 | 0.052 | 0.120 | Cuneus |
| 0.108 | 0.062 | 0.110 | 0.073 | Entorhinal cortex |
| 0.070 | 0.088 | 0.054 | 0.083 | Frontal pole |
| 0.083 | 0.065 | 0.119 | 0.073 | Fusiform gyrus |
| 0.094 | 0.074 | 0.098 | 0.074 | Inferior parietal gyrus |
| 0.110 | 0.085 | 0.111 | 0.071 | Inferior temporal gyrus |
| 0.076 | 0.052 | 0.086 | 0.052 | Insula |
| 0.072 | 0.077 | 0.082 | 0.093 | Isthmus cingulate cortex |
| 0.067 | 0.094 | 0.081 | 0.082 | Lateral occipital gyrus |
| 0.081 | 0.070 | 0.082 | 0.062 | Lateral orbitofrontal cortex |
| 0.075 | 0.076 | 0.062 | 0.087 | Lingual gyrus |
| 0.070 | 0.079 | 0.066 | 0.059 | Medial orbitofrontal cortex |
| 0.091 | 0.065 | 0.097 | 0.056 | Middle temporal gyrus |
| 0.065 | 0.089 | 0.074 | 0.060 | Paracentral gyrus |
| 0.089 | 0.077 | 0.098 | 0.074 | Parahippocampal gyrus |
| 0.091 | 0.064 | 0.071 | 0.068 | Pars opercularis of the inferior frontal gyrus |
| 0.113 | 0.071 | 0.089 | 0.070 | Pars orbitalis of the inferior frontal gyrus |
| 0.059 | 0.083 | 0.071 | 0.089 | Pars triangularis of the inferior frontal gyrus |
| 0.048 | 0.101 | 0.057 | 0.121 | Pericalcarine gyrus |
| 0.064 | 0.094 | 0.061 | 0.087 | Postcentral gyrus |
| 0.072 | 0.082 | 0.084 | 0.078 | Posterior cingulate cortex |
| 0.100 | 0.058 | 0.090 | 0.072 | Precentral gyrus |
| 0.088 | 0.067 | 0.091 | 0.074 | Precuneus |
| 0.073 | 0.095 | 0.075 | 0.091 | Rostral anterior cingulate cortex |
| 0.078 | 0.083 | 0.067 | 0.080 | Rostral middle frontal gyrus |
| 0.084 | 0.079 | 0.070 | 0.071 | Superior frontal gyrus |
| 0.086 | 0.077 | 0.076 | 0.092 | Superior parietal gyrus |
| 0.086 | 0.083 | 0.103 | 0.068 | Superior temporal gyrus |
| 0.072 | 0.067 | 0.088 | 0.078 | Supramarginal gyrus |
| 0.135 | 0.067 | 0.110 | 0.060 | Temporal pole |
| 0.089 | 0.077 | 0.087 | 0.078 | Transversetemporal gyrus |
| 0.083 | 0.071 | 0.085 | 0.076 | Average cortical thickness |

Percentage of extreme deviations in one region, measured against the number of individuals with at least one extreme deviation.

**S1.9: Percentages of train healthy controls, test healthy controls and individuals with depression with an extreme deviation (positive deviation: z > 1.96; negative deviation z< -1.96), breakdown region-wise, per set**

| Healthy controls, training set | | Healthy controls, test set | | MDD, test set | | |
|---|---|---|---|---|---|---|
| Neg. dev. | Pos. dev. | Neg. dev. | Pos. dev. | Neg. dev. | Pos. dev. | Region of Interest |
| 0.027 | 0.026 | 0.027 | 0.030 | 0.032 | 0.022 | Banks of the Superior Temporal Sulcus |
| 0.021 | 0.024 | 0.024 | 0.030 | 0.023 | 0.027 | Caudal anterior cingulate cortex |
| 0.025 | 0.022 | 0.030 | 0.025 | 0.024 | 0.023 | Caudal middle frontal gyrus |
| 0.021 | 0.030 | 0.016 | 0.037 | 0.019 | 0.037 | Cuneus |
| 0.026 | 0.021 | 0.032 | 0.020 | 0.038 | 0.022 | Entorhinal cortex |
| 0.020 | 0.026 | 0.021 | 0.028 | 0.018 | 0.025 | Frontal Pole |
| 0.027 | 0.025 | 0.025 | 0.021 | 0.040 | 0.022 | Fusiform gyrus |
| 0.024 | 0.022 | 0.028 | 0.024 | 0.033 | 0.023 | Inferior parietal gyrus |
| 0.023 | 0.022 | 0.033 | 0.027 | 0.038 | 0.021 | Inferior temporal gyrus |
| 0.024 | 0.019 | 0.023 | 0.017 | 0.029 | 0.016 | Insula |
| 0.017 | 0.029 | 0.021 | 0.025 | 0.028 | 0.028 | Isthmus cingulate cortex |
| 0.019 | 0.025 | 0.020 | 0.030 | 0.027 | 0.026 | Lateral occipital gyrus |
| 0.024 | 0.027 | 0.024 | 0.022 | 0.028 | 0.019 | Lateral orbitofrontal cortex. |
| 0.019 | 0.030 | 0.022 | 0.024 | 0.021 | 0.027 | Lingual gyrus |
| 0.023 | 0.026 | 0.021 | 0.025 | 0.022 | 0.018 | Medial orbitofrontal cortex. |
| 0.024 | 0.022 | 0.027 | 0.021 | 0.033 | 0.017 | Middle temporal cortex. |
| 0.026 | 0.021 | 0.019 | 0.028 | 0.026 | 0.019 | Paracentral gyrus |
| 0.024 | 0.021 | 0.026 | 0.025 | 0.034 | 0.022 | Parahippocampal gyrus |
| 0.025 | 0.027 | 0.027 | 0.020 | 0.024 | 0.021 | Pars opercularis of the inferior frontal gyrus |
| 0.028 | 0.025 | 0.034 | 0.023 | 0.030 | 0.021 | Pars orbitalis of the inferior frontal gyrus |
| 0.023 | 0.025 | 0.018 | 0.026 | 0.025 | 0.027 | Pars triangularis of the inferior frontal gyrus |
| 0.016 | 0.031 | 0.015 | 0.033 | 0.020 | 0.037 | Pericalcarine gyrus |
| 0.020 | 0.025 | 0.019 | 0.030 | 0.021 | 0.028 | Postcentral gyrus |
| 0.022 | 0.027 | 0.021 | 0.026 | 0.029 | 0.023 | Posterior cingulate cortex |
| 0.031 | 0.019 | 0.030 | 0.018 | 0.032 | 0.022 | Precentral gyrus |
| 0.025 | 0.023 | 0.026 | 0.021 | 0.031 | 0.022 | precuneus |
| 0.025 | 0.024 | 0.022 | 0.030 | 0.026 | 0.027 | Rostral anterior cingulate cortex |
| 0.021 | 0.025 | 0.023 | 0.026 | 0.023 | 0.024 | Rostral middle frontal gyrus |
| 0.026 | 0.023 | 0.025 | 0.025 | 0.024 | 0.022 | Superior frontal gyrus |
| 0.025 | 0.024 | 0.025 | 0.025 | 0.026 | 0.029 | Superior parietal gyrus |
| 0.025 | 0.024 | 0.025 | 0.026 | 0.035 | 0.021 | Superior temporal gyrus |
| 0.028 | 0.022 | 0.021 | 0.021 | 0.030 | 0.023 | Supramarginal gyrus |
| 0.036 | 0.021 | 0.040 | 0.021 | 0.038 | 0.018 | Temporal pole |
| 0.021 | 0.027 | 0.027 | 0.025 | 0.030 | 0.024 | Transversetemporal gyrus |
| 0.026 | 0.022 | 0.025 | 0.023 | 0.029 | 0.024 | Average cortical thickness |

positive deviation: z > 1.96; negative deviation z< -1.96, breakdown region-wise. Pos dev: positive deviations. Neg. dev: negative deviations

#### S1.10: Model fit in Stan: Correlation coefficient /rho between observed and predicted value per set, region-wise breakdown.

| Region of interest | HC Training set | HC Test set | MDD set |
|---|---|---|---|
| Banks of the Superior Temporal Sulcus | 0.62 | 0.59 | 0.58 |
| Caudal anterior cingulate gyrus | 0.58 | 0.54 | 0.54 |
| Caudal middle frontal cortex | 0.76 | 0.74 | 0.68 |
| Cuneus | 0.79 | 0.79 | 0.69 |
| Entorhinal cortex | 0.65 | 0.65 | 0.51 |
| Frontal Pole | 0.56 | 0.57 | 0.55 |
| Fusiform gyrus | 0.72 | 0.72 | 0.72 |
| Inferior parietal gyrus | 0.77 | 0.77 | 0.69 |
| Inferior temporal gyrus | 0.78 | 0.77 | 0.75 |
| Insula | 0.64 | 0.62 | 0.65 |
| Isthmus cingulate cortex | 0.69 | 0.70 | 0.68 |
| Lateral occipital gyrus | 0.76 | 0.77 | 0.67 |
| Lateral orbitofrontal cortex. | 0.62 | 0.59 | 0.59 |
| Lingual gyrus | 0.71 | 0.71 | 0.62 |
| Medial orbitofrontal cortex. | 0.66 | 0.65 | 0.65 |
| Middle temporal gyrus. | 0.69 | 0.68 | 0.64 |
| Paracentral gyrus | 0.87 | 0.88 | 0.79 |
| Parahippocampal gyrus | 0.43 | 0.44 | 0.43 |
| Pars opercularis of the inferior frontal gyrus. | 0.74 | 0.73 | 0.66 |
| Pars orbitalis of the inferior frontal gyrus | 0.59 | 0.56 | 0.50 |
| Pars triangularis of the inferior frontal gyrus | 0.72 | 0.73 | 0.66 |
| Pericalcarine gyrus. | 0.82 | 0.82 | 0.74 |
| Postcentral gyrus | 0.80 | 0.81 | 0.70 |
| Posterior cingulate cortex | 0.74 | 0.74 | 0.73 |
| Precentral gyrus | 0.85 | 0.85 | 0.75 |
| precuneus | 0.80 | 0.79 | 0.72 |
| Rostral anterior cingulate cortex | 0.58 | 0.52 | 0.53 |
| Rostral middle frontal gyrus | 0.74 | 0.72 | 0.68 |
| Superior frontal gyrus | 0.76 | 0.75 | 0.70 |
| Superior parietal gyrus | 0.79 | 0.79 | 0.71 |
| Superior temporal gyrus | 0.66 | 0.64 | 0.57 |
| Supramarginal gyrus | 0.79 | 0.78 | 0.70 |
| Temporal pole | 0.66 | 0.63 | 0.50 |
| Transversetemporal gyrus | 0.59 | 0.57 | 0.51 |
| Average cortical thickness | 0.79 | 0.79 | 0.70 |

#### S1.11: Model fit in Stan: Standardized root mean squared errors (SRMSE) per set, region-wise breakdown.

| Region of interest | HC Training set | HC Test set | MDD |
|---|---|---|---|
| Banks of the Superior Temporal Sulcus | 0.77 | 0.81 | 0.79 |
| Caudal anterior cingulate cortex | 0.81 | 0.83 | 0.83 |
| Caudal middle frontal gyrus | 0.65 | 0.67 | 0.66 |
| Cuneus | 0.61 | 0.61 | 0.63 |
| Entorhinal cortex | 0.75 | 0.74 | 0.79 |
| Frontal pole | 0.83 | 0.82 | 0.82 |
| Fusiform gyrus | 0.70 | 0.70 | 0.74 |
| Inferior parietal gyrus | 0.63 | 0.64 | 0.66 |
| Inferior temporal gyrus | 0.62 | 0.65 | 0.65 |
| Insula | 0.77 | 0.76 | 0.77 |
| Isthmus cingulate cortex | 0.72 | 0.72 | 0.76 |
| Lateral occipital gyrus | 0.65 | 0.65 | 0.67 |
| Lateral orbitofrontal cortex | 0.79 | 0.77 | 0.79 |
| Lingual gyrus | 0.70 | 0.69 | 0.70 |
| Medial orbitofrontal cortex | 0.75 | 0.74 | 0.72 |
| Middle temporal gyrus | 0.71 | 0.73 | 0.73 |
| Paracentral gyrus | 0.50 | 0.49 | 0.50 |
| Parahippocampal gyrus | 0.90 | 0.92 | 0.93 |
| Pars opercularis of the inferior frontal gyrus | 0.67 | 0.67 | 0.69 |
| Pars orbitalis of the inferior frontal gyrus | 0.81 | 0.83 | 0.83 |
| Pars triangularis of the inferior frontal gyrus | 0.70 | 0.68 | 0.70 |
| Pericalcarine gyrus | 0.58 | 0.57 | 0.60 |
| Postcentral gyrus | 0.60 | 0.59 | 0.61 |
| Posterior cingulate cortex | 0.67 | 0.66 | 0.68 |
| Precentral gyrus | 0.52 | 0.53 | 0.54 |
| Precuneus | 0.60 | 0.61 | 0.63 |
| Rostral anterior cingulate cortex | 0.81 | 0.83 | 0.84 |
| Rostral middle frontal gyrus | 0.68 | 0.69 | 0.67 |
| Superior frontal gyrus | 0.64 | 0.67 | 0.66 |
| Superior parietal gyrus | 0.61 | 0.61 | 0.64 |
| Superior temporal gyrus | 0.74 | 0.75 | 0.77 |
| Supramarginal gyrus | 0.61 | 0.62 | 0.63 |
| Temporal pole | 0.75 | 0.77 | 0.76 |
| Transversetemporal gyrus | 0.81 | 0.81 | 0.83 |
| Average cortical thickness | 0.61 | 0.62 | 0.63 |

#### S1.12: Model fit in Stan: Explained variance of the predictions per set, region-wise breakdown.

| Region of interest | HC Training set | HC Test set | MDD |
|---|---|---|---|
| Banks of the Superior Temporal Sulcus | 0.38 | 0.35 | 0.34 |
| Caudal anterior cingulate cortex | 0.34 | 0.29 | 0.29 |
| Caudal middle frontal gyrus | 0.57 | 0.55 | 0.46 |
| Cuneus | 0.62 | 0.63 | 0.48 |
| Entorhinal cortex | 0.42 | 0.42 | 0.25 |
| Frontal pole | 0.31 | 0.32 | 0.30 |
| Fusiform gyrus | 0.51 | 0.52 | 0.52 |
| Inferior parietal gyrus | 0.60 | 0.59 | 0.47 |
| Inferior temporal gyrus | 0.61 | 0.60 | 0.57 |
| Insula | 0.41 | 0.38 | 0.42 |
| Isthmus cingulate cortex | 0.48 | 0.49 | 0.45 |
| Lateral occipital gyrus | 0.58 | 0.59 | 0.45 |
| Lateral orbitofrontal cortex | 0.38 | 0.34 | 0.34 |
| Lingual gyrus | 0.51 | 0.51 | 0.38 |
| Medial orbitofrontal cortex | 0.44 | 0.43 | 0.42 |
| Middle temporal gyrus | 0.48 | 0.47 | 0.41 |
| Paracentral gyrus | 0.75 | 0.77 | 0.62 |
| Parahippocampal gyrus | 0.18 | 0.19 | 0.18 |
| Pars opercularis of the inferior frontal gyrus | 0.55 | 0.54 | 0.43 |
| Pars orbitalis of the inferior frontal gyrus | 0.35 | 0.31 | 0.23 |
| Pars triangularis of the inferior frontal gyrus | 0.52 | 0.53 | 0.43 |
| Pericalcarine gyrus | 0.66 | 0.67 | 0.55 |
| Postcentral gyrus | 0.64 | 0.65 | 0.49 |
| Posterior cingulate cortex | 0.55 | 0.55 | 0.53 |
| Precentral gyrus | 0.73 | 0.72 | 0.55 |
| Precuneus | 0.64 | 0.63 | 0.52 |
| Rostral anterior cingulate cortex | 0.33 | 0.27 | 0.28 |
| Rostral middle frontal gyrus | 0.54 | 0.52 | 0.46 |
| Superior frontal gyrus | 0.58 | 0.56 | 0.49 |
| Superior parietal gyrus | 0.63 | 0.63 | 0.50 |
| Superior temporal gyrus | 0.44 | 0.41 | 0.32 |
| Supramarginal gyrus | 0.62 | 0.61 | 0.48 |
| Temporal pole | 0.43 | 0.40 | 0.24 |
| Transversetemporal gyrus | 0.35 | 0.33 | 0.26 |
| Average cortical thickness | 0.63 | 0.62 | 0.48 |

#### S1.13: Model fit in Stan: Mean standardized log loss per set, region-wise breakdown.

| Region of interest | HC Test set | MDD set |
|---|---|---|
| Banks of the Superior Temporal Sulcus | -0.10 | -0.12 |
| Caudal anterior cingulate cortex | 0.01 | 0.10 |
| Caudal middle frontal gyrus | -0.39 | -0.32 |
| Cuneus | -0.45 | -0.35 |
| Entorhinal cortex | -0.22 | 0.11 |
| Frontal pole | 0.05 | 0.04 |
| Fusiform gyrus | -0.36 | -0.36 |
| Inferior parietal gyrus | -0.42 | -0.32 |
| Inferior temporal gyrus | -0.43 | -0.40 |
| Insula | -0.19 | -0.23 |
| Isthmus cingulate cortex | -0.33 | -0.27 |
| Lateral occipital gyrus | -0.42 | -0.32 |
| Lateral orbitofrontal cortex | -0.13 | -0.15 |
| Lingual gyrus | -0.36 | -0.21 |
| Medial orbitofrontal cortex | -0.27 | -0.26 |
| Middle temporal gyrus | -0.31 | -0.23 |
| Paracentral gyrus | -0.50 | -0.46 |
| Parahippocampal gyrus | 1.22 | 0.97 |
| Pars opercularis of the inferior frontal gyrus | -0.38 | -0.28 |
| Pars orbitalis of the inferior frontal gyrus | 0.00 | 0.04 |
| Pars triangularis of the inferior frontal gyrus | -0.37 | -0.28 |
| Pericalcarine gyrus | -0.46 | -0.40 |
| Postcentral gyrus | -0.45 | -0.36 |
| Posterior cingulate cortex | -0.39 | -0.38 |
| Precentral gyrus | -0.47 | -0.41 |
| Precuneus | -0.44 | -0.37 |
| Rostral anterior cingulate cortex | 0.05 | 0.23 |
| Rostral middle frontal gyrus | -0.37 | -0.31 |
| Superior frontal gyrus | -0.40 | -0.35 |
| Superior parietal gyrus | -0.44 | -0.35 |
| Superior temporal gyrus | -0.24 | -0.08 |
| Supramarginal gyrus | -0.43 | -0.34 |
| Temporal pole | -0.20 | 0.01 |
| Transversetemporal gyrus | -0.05 | 0.32 |
| Average cortical thickness | -0.43 | -0.34 |

### S1.14 Average z-score predicting clinical variables.

**Categorical variables: logistic regression**

| **adjusted p values** | odds (RR) | variables | baseline (odds=1) |
|---|---|---|---|
| **<0.001** | 0.83 | Dx | HC |
| **<0.001** | 0.78 | First | HC |
| **<0.001** | 0.84 | Recur | HC |
| **0.19** | 1.08 | First | Recur |
| **<0.001** | 0.78 | Acu | HC |
| **0.16** | 0.88 | Rem | HC |
| **0.19** | 0.89 | Acu | Rem |
| **<0.001** | 0.77 | AD | HC |
| **0.03** | 0.88 | noAD | HC |
| **0.07** | 0.90 | noAD | AD |
| **<0.001** | 0.78 | EAO | HC |
| **<0.001** | 0.84 | LAO | HC |
| **0.19** | 1.08 | EAO | LAO |

**Continuous variables - linear regression**

| **adjusted p values** | beta | scores |
|---|---|---|
| **0.13** | -0.48 | HDRS |
| **0.05** | -0.99 | BDI |
| **<0.001** | -0.99 | BMI |
| **0.24** | -0.72 | CTQ |

Dx: depressed. HC: healthy control. First: First episode depression. Recur: Recurrent episode depression. Acu: Acutely depressed. Rem: Remitted. AD: Anti-depressant using patient at the time of the scan. noAD: non-Anti-depressant using patient at the time of the scan. EAO: Early onset depression (=< 21 Years). LAO: Late onset depression (> 21 Years). HDRS: Hamilton Depression Rating Scale. BDI: Beck Depression Inventory. BMI: Body Mass Index. CTQ: Childhood Trauma Questionnaire.

#### S1.15 Positive load predicting clinical variables.

**Categorical variables: logistic regression**

| adjusted p values | odds (RR) | variables | Baseline (odds=1) |
|---|---|---|---|
| **0.49** | 0.99 | Dx | HC |
| **0.49** | 0.98 | First | HC |
| **0.66** | 0.99 | Recur | HC |
| **0.74** | 1.01 | First | Recur |
| **0.34** | 0.98 | Acu | HC |
| **0.34** | 1.03 | Rem | HC |
| **0.10** | 0.95 | Acu | Rem |
| **0.10** | 0.97 | AD | HC |
| **0.71** | 1.01 | noAD | HC |
| **0.08** | 0.96 | noAD | AD |
| **0.30** | 0.97 | EAO | HC |
| **0.36** | 0.98 | LAO | HC |
| **0.74** | 1.01 | EAO | LAO |

**Continuous variables - linear regression**

| adjusted p values | beta | variables |
|---|---|---|
| **0.05** | -0.24 | HDRS |
| **0.29** | -0.23 | BDI |
| **0.05** | -0.19 | BMI |
| **0.77** | -0.06 | CTQ |

Dx: depressed. HC: healthy control. First: First episode depression. Recur: Recurrent episode depression. Acu: Acutely depressed. Rem: Remitted. AD: Anti-depressant using patient at the time of the scan. noAD: non-Anti-depressant using patient at the time of the scan. EAO: Early onset depression (=< 21 Years). LAO: Late onset depression (> 21 Years). HDRS: Hamilton Depression Rating Scale. BDI: Beck Depression Inventory. BMI: Body Mass Index. CTQ: Childhood Trauma Questionnaire.

### S1.16 Negative load predicting clinical variables.

**Categorical variables: logistic regression**

| adjusted p values | Odds (RR) | variables | baseline (odds=1) |
|---|---|---|---|
| **0.11** | 1.02 | Dx | HC |
| **0.11** | 1.03 | First | HC |
| **0.33** | 1.02 | Recur | HC |
| **0.36** | 0.98 | First | Recur |
| **0.15** | 1.02 | Acu | HC |
| **0.11** | 1.03 | Rem | HC |
| **0.36** | 0.98 | Acu | Rem |
| **0.56** | 1.01 | AD | HC |
| **0.05** | 1.03 | noAD | HC |
| **0.11** | 0.97 | noAD | AD |
| **0.05** | 1.04 | EAO | HC |
| **0.83** | 1.00 | LAO | HC |
| **0.05** | 0.96 | EAO | LAO |

**Continuous variables: linear regression**

| adjusted p values | beta | variables |
|---|---|---|
| **0.11** | -0.14 | HDRS |
| **0.46** | 0.11 | BDI |
| **0.05** | 0.16 | BMI |
| **0.59** | 0.08 | CTQ |

Dx: depressed. HC: healthy control. First: First episode depression. Recur: Recurrent episode depression. Acu: Acutely depressed. Rem: Remitted. AD: Anti-depressant using patient at the time of the scan. noAD: non-Anti-depressant using patient at the time of the scan. EAO: Early onset depression (=< 21 Years). LAO: Late onset depression (> 21 Years). HDRS: Hamilton Depression Rating Scale. BDI: Beck Depression Inventory. BMI: Body Mass Index. CTQ: Childhood Trauma Questionnaire.

#### S1.17 Positive extremity predicting clinical variables.

**Categorical variables: logistic regression**

| adjusted p values | Odds (RR) | Clinical variables | baseline (odds=1) |
|---|---|---|---|
| **0.11** | 0.932 | Dx | HC |
| **0.26** | 0.936 | First | HC |
| **0.15** | 0.929 | Recur | HC |
| **0.89** | 0.994 | First | Recur |
| **0.11** | 0.920 | Acu | HC |
| **0.26** | 1.095 | Rem | HC |
| **0.04** | 0.847 | Acu | Rem |
| **0.01** | 0.871 | AD | HC |
| **0.89** | 0.988 | noAD | HC |
| **0.03** | 0.888 | noAD | AD |
| **0.23** | 0.930 | EAO | HC |
| **0.02** | 0.886 | LAO | HC |
| **0.39** | 0.954 | EAO | LAO |

**Continuous variables - linear regression**

| adjusted p values | beta | variables |
|---|---|---|
| **0.04** | -0.55 | HDRS |
| **0.26** | -0.49 | BDI |
| **0.01** | -0.66 | BMI |
| **0.89** | -0.11 | CTQ |

Dx: depressed. HC: healthy control. First: First episode depression. Recur: Recurrent episode depression. Acu: Acutely depressed. Rem: Remitted. AD: Anti-depressant using patient at the time of the scan. noAD: non-Anti-depressant using patient at the time of the scan. EAO: Early onset depression (=< 21 Years). LAO: Late onset depression (> 21 Years). HDRS: Hamilton Depression Rating Scale. BDI: Beck Depression Inventory. BMI: Body Mass Index. CTQ: Childhood Trauma Questionnaire.

#### S1.18 Negative extremity predicting clinical variables.

**Categorical variables: logistic regression**

| adjusted p values | Odds (RR) | variables | Baseline (odds=1) |
|---|---|---|---|
| **<0.001** | 1.14 | Dx | HC |
| **<0.001** | 1.20 | First | HC |
| **<0.001** | 1.15 | Recur | HC |
| **0.47** | 0.96 | First | Recur |
| **<0.001** | 1.22 | Acu | HC |
| **<0.001** | 1.27 | Rem | HC |
| **0.42** | 0.94 | Acu | Rem |
| **<0.001** | 1.15 | AD | HC |
| **0.01** | 1.13 | noAD | HC |
| **0.90** | 1.01 | noAD | AD |
| **<0.001** | 1.21 | EAO | HC |
| **0.12** | 1.07 | LAO | HC |
| **0.02** | 0.89 | EAO | LAO |

**Continuous variables - linear regression**

| adjusted p values | beta | variables |
|---|---|---|
| **0.95** | -0.02 | HDRS |
| **0.05** | 0.82 | BDI |
| **0.03** | 0.49 | BMI |
| **0.24** | 0.69 | CTQ |

Dx: depressed. HC: healthy control. First: First episode depression. Recur: Recurrent episode depression. Acu: Acutely depressed. Rem: Remitted. AD: Anti-depressant using patient at the time of the scan. noAD: non-Anti-depressant using patient at the time of the scan. EAO: Early onset depression (=< 21 Years). LAO: Late onset depression (> 21 Years). HDRS: Hamilton Depression Rating Scale. BDI: Beck Depression Inventory. BMI: Body Mass Index. CTQ: Childhood Trauma Questionnaire.

#### S1. 19. Comparison of impact of imputation of test MDD set by train healthy controls vs, by test MDD set, corrected for multiple comparisons.

| ROIs | Mean, imputed on HC | Mean, imputed on MDD | P value of difference |
|---|---|---|---|
| Banks of the Superior Temporal Sulcus | -0.048 | -0.0103 | 0.1648 |
| Caudal anterior cingulate cortex | -0.072 | -0.0001 | 0.0058 |
| Caudal middle frontal gyrus | 0.008 | 0.0014 | 0.79 |
| Cuneus | 0.1537 | 0.002 | **> 0.0001** |
| Entorhinal cortex | -0.0358 | -0.0018 | 0.1982 |
| Fusiform gyrus | -0.0131 | -0.003 | 0.7759 |
| Inferior parietal gyrus | 0.0138 | 0.0011 | 0.74 |
| Inferior temporal gyrus | -0.0319 | -0.0215 | 0.7759 |
| Isthmus cingulate cortex | -0.0411 | 0.0006 | 0.1407 |
| Lateral occipital gyrus | 0.1992 | -0.0006 | **> 0.0001** |
| Lateral orbitofrontal cortex | -0.0097 | -0.0008 | 0.7759 |
| Lingual gyrus | 0.0789 | -0.0005 | **0.0012** |
| Medial orbitofrontal cortex | -0.0837 | 0.0008 | 0.0009 |
| Middle temporal gyrus | -0.0475 | -0.0068 | 0.1381 |
| Parahippocampal gyrus | -0.0303 | 0.0003 | 0.2885 |
| Paracentral gyrus | 0.062 | -0.0005 | **0.0088** |
| Pars opercularis of the inferior frontal gyrus | -0.0585 | 0.0003 | **0.0199** |
| Pars orbitalis of the inferior frontal gyrus | -0.0389 | 0.001 | 0.1407 |
| Pars triangularis of the inferior frontal gyrus | -0.0104 | 0 | 0.7759 |
| Pericalcarine gyrus | 0.1436 | -0.0015 | **> 0.0001** |
| Postcentral gyrus | 0.0899 | -0.0012 | **0.0002** |
| Posterior cingulate cortex | -0.0244 | 0 | 0.411 |
| Precentral gyrus | 0.0805 | 0.0027 | **0.001** |
| Precuneus | 0.0066 | -0.0012 | 0.7759 |
| Rostral anterior cingulate cortex | -0.1329 | -0.0005 | **> 0.0001** |
| Rostral middle frontal gyrus | -0.0154 | 0.001 | 0.6243 |
| Superior frontal gyrus | -0.0826 | 0.0019 | **0.0009** |
| Superior parietal gyrus | 0.069 | 0.0011 | **0.0061** |
| Superior temporal gyrus | -0.0794 | -0.0074 | **0.0044** |
| Supramarginal gyrus | 0.0088 | 0.0013 | 0.7759 |
| Frontal pole | 0.0462 | -0.0002 | 0.0964 |
| Temporal pole | 0.0268 | -0.0034 | 0.2569 |
| Transversetemporal gyrus | -0.0074 | -0.0024 | 0.8265 |
| Insula | -0.1573 | 0.0001 | **> 0.0001** |
| Average cortical thickness | 0.0437 | 0 | 0.0964 |

S1.19 Comparison of imputation of MDD test set by HC training set vs. by itself. MDD: Major Depressive Disorder. HC: healthy controls
